# Matching Habitat Choice and the Evolution of a Species’ Range

**DOI:** 10.1101/2024.06.25.600700

**Authors:** Farshad Shirani, Judith R. Miller

## Abstract

Natural selection is not the only mechanism that promotes adaptation of an organism to its environment. Another mechanism is matching habitat choice, in which individuals sense and disperse toward habitat best suited to their phenotype. This can in principle facilitate rapid adaptation, enhance range expansion, and promote genetic differentiation, reproductive isolation, and speciation. However, empirical evidence that confirms the evolution of matching habitat choice in nature is limited. Here we obtain theoretical evidence that phenotype-optimal dispersal, a particular form of matching habitat choice, is likely to evolve only in the presence of a steep environmental gradient. Such a gradient may be steeper than the gradient the majority of species typically experience in nature, adding to the collection of possible explanations for the scarcity of evidence for matching habitat choice. We draw this conclusion from numerical solutions of a system of deterministic partial differential equations for a population’s density along with the mean and variance of a fitness-related quantitative phenotypic trait such as body size. In steep gradients, we find that phenotype-optimal dispersal facilitates rapid adaptation on single-generation time scales, reduces within-population trait variation, increases range expansion speed, and enhances the chance of survival in rapidly changing environments. Moreover, it creates a directed gene flow that compensates for the maladaptive core-to-edge effects of random gene flow caused by random movements. These results suggest that adaptive gene flow to range margins, together with substantially reduced trait variation at central populations, may be hallmarks of phenotype-optimal dispersal in natural populations. Further, slowly-growing species under strong natural selection may particularly benefit from evolving phenotype-optimal dispersal.

## 1. Introduction

A species’ range evolves through complex eco-evolutionary processes, the understanding of which is crucial for predicting species’ response to climate change, informing conservation efforts for preserving biodiversity, and developing strategies for controlling invasive species (Angert et al., 2020; Bridle and Vines, 2007; Brown et al., 1996; Case et al., 2005; Duckworth and Badyaev, 2007; Fronhofer and Altermatt, 2015; Gaston et al., 2003; Godsoe et al., 2017; Haddad et al., 2015; Hoffmann and Blows, 1994; Holt and Keitt, 2005; Keitt et al., 2001; Kirkpatrick and Barton, 1997; Louthan et al., 2015; MacArthur, 1972; Miller et al., 2020; Ponchon and Travis, 2022; Pulliam, 2000; Rafajlović et al., 2022; Sexton et al., 2009; Shirani and Miller, 2022). Dispersal is one of the key ecological factors involved in determining a species’ range. Since dispersal creates gene flow, it affects the spatial dynamics of genetic variance and local adaptation, as well as population dynamic measures such as range expansion rates (Baguette et al., 2013; Bonte et al., 2012; Bowler and Benton, 2005; Gomulkiewicz et al., 1999; Holt and Gomulkiewicz, 1997; Jacob et al., 2015; Kawecki and Ebert, 2004; Kirkpatrick and Barton, 1997; Okubo and Levin, 2001; Pellerin et al., 2019; Ronce, 2007; Slatkin, 1985, 1987).

The simplest types of dispersal, including the archetype of diffusive spread, are in theory independent of individual, population and environmental characteristics (Lowe and McPeek, 2014). In general, random dispersal of individuals to other habitat locations irrespective of their genotypes imposes a *migration load* on the population’s mean fitness (Bolnick and Otto, 2013; Bonte et al., 2012; Edelaar and Bolnick, 2012; Lenormand, 2002). By contrast, a population’s fitness increases, most generally, when its individuals evolve adaptive dispersal strategies based on their environment structure, their own characteristics, and their intra- and interspecific interactions (Baguette et al., 2013; Baines et al., 2019; Bonte et al., 2012; Bowler and Benton, 2005; Clobert et al., 2009; Cote et al., 2017; Edelaar and Bolnick, 2012; Holt, 1985, 1987; Jaenike and Holt, 1991; Levins, 1963; Lustenhouwer et al., 2023; Rausher, 1984; Ronce, 2007; Saastamoinen et al., 2018; Travis et al., 2012). Such convoluted evolution of dispersal—as a complex multi-dimensional phenotype comprising of different components (propensity to disperse, distance and direction of movements, and settlement choice (Saastamoinen et al., 2018))—is driven by the balance between the cost incurred at each stage of dispersal and the overall benefit acquired from dispersal (Bonte et al., 2012; Bowler and Benton, 2005; Cote et al., 2017; Garant et al., 2007; Rosenzweig, 1981).

Empirical evidence in a variety of species confirms that individuals often bias their dispersal towards preferred habitat locations (Armsworth and Roughgarden, 2008; Bolnick and Otto, 2013; Cote et al., 2017; Fretwell, 1972; Holt, 1987; Jacob et al., 2015; Lowe and McPeek, 2014; Pellerin et al., 2019; Rice and Salt, 1990; Ronce, 2007). To carry out such fitness-associated dispersal, individuals of such species must perceive their internal phenotypic traits, and must efficiently orient their movements by actively using different sources of information, such as abiotic cues, landscape landmarks, and presence and behavior of conspecifics. For instance, nocturnal snakes can sense the temperature and physical structure of rocks to move to thermally preferable habitats (Bowler and Benton, 2005; Clobert et al., 2009; Ponchon and Travis, 2022).

Here, we propose and numerically study a deterministic model of a species’ range evolution under a fairly idealized habitat-matching strategy. The dispersal strategy that we study has been hypothesized to have substantial and distinctive effects on species evolutionary range dynamics (Edelaar et al., 2008). We demonstrate such effects through numerical simulations and identify conditions under which this adaptive dispersal strategy will be sufficiently beneficial to evolve.

### 1.1 Matching Habitat Choice: An Adaptive Dispersal Strategy

In the most idealized version of adaptive dispersal, individuals can perceive all components of their absolute fitness and are able to move freely. Therefore, they can climb local fitness gradients and achieve their maximum expected fitness (Armsworth, 2009; Armsworth and Roughgarden, 2005, 2008; Fretwell, 1972; Holt, 1985, 1987; Ravigné et al., 2009; Rosenzweig, 1981; Ruxton and Rohani, 1999). However, acquiring information about all major components of fitness is infeasible. In a still idealized but more realistic conception, dispersing individuals move toward habitat that is best for their fitness-related phenotypes. For example, medium ground finches with deeper bills, which can crack larger seeds, tend to settle in areas richer in large-seeded plants to increase their food intake (Edelaar and Bolnick, 2019). Therefore, with this type of phenotype-dependent adaptive dispersal—which is often referred to as *matching habitat choice* (Edelaar et al., 2008; Ravigné et al., 2004)—the expected performance of the individuals is maximized.

Although the maximized performance under matching habitat choice is also likely to lead to a considerable increase in the individuals’ fitness (Edelaar and Bolnick, 2019)—due to the presumed correlation between individuals’ phenotype and fitness—it is worth clarifying the difference between matching habitat choice (as phenotype-optimal dispersal) and fitness-optimal dispersal. Matching habitat choice is a phenotype-environment matching strategy that can involve following an environmental gradient in optimum phenotype. This is not necessarily the same as following the gradient in optimal (maximum) fitness. In general, as we described above, matching habitat choice is often conceptualized as a “performance”-maximizing but not necessarily a “fitness”-maximizing strategy. For example, grasshoppers choosing a habitat based on a match between their color and the thermal radiation of the habitat are not necessarily maximizing their fitness by doing so. In fact, many factors affecting population fitness, such as resource availability, mating success, fecundity, and importantly, competition do not directly contribute to adopting a matching habitat choice strategy.

Although strong evidence for matching habitat choice in nature is still limited, a growing number of experimental and empirical studies have identified it in diverse taxa. For instance, using microcosms of a ciliate species which shows genetic variability in performance along a thermal gradient, Jacob et al. (2017) have experimentally shown that local adaptation to the upper margin of the species’ thermal niche is favored by dispersal with matching habitat choice, whereas it is hindered under random dispersal. Similarly, in a semi-natural warming experiment with a model species of reptiles, Bestion et al. (2015) have shown that individuals disperse to warmer or cooler habitats based on their preferred temperature. In another study, Camacho et al. (2020) performed a color manipulation experiment on grasshoppers to test for camouflage-based background matching in an urban mosaic of dark and pale pavements. They found that black-painted grasshoppers mostly chose to settle in dark asphalt, whereas white-painted ones chose pale pavement—the same matching substrate choice behavior observed naturally in unmanipulated morphs. A similar study by Karpestam et al. (2012) on color-manipulated pygmy grasshoppers over a solar radiation mosaic demonstrated that black-painted individuals tended to reside in habitats with less radiation, and white-painted females had more hatchlings than black-painted ones in increased radiation treatments. Color-dependent habitat choice has also been identified in dark and pale barn owls (Dreiss et al., 2012). For a species of nomadic crossbill bird, matching habitat choice has been proposed as a contributor to rapid diversification of ecotypes; this hypothesis was supported by mark-recapture data (Benkman, 2017).

### 1.2 Eco-evolutionary Impacts of Matching Habitat Choice

With matching habitat choice, individuals sort themselves across the environment to minimize their phenotype-environment mismatch. This means that, matching habitat choice is in fact an adaptation mechanism that operates at the individual level and, in principle, results in rapid population adaptation on *within-generation* timescales—that is, a strong match between mean population phenotype and environment optimum phenotype can occur within one generation time. This mode of rapid adaptation is essentially different from adaptation by natural selection, which operates only at the population level. Although, in essence, the process of adaptation by natural selection also takes place within generations, plausible levels of natural selection often require many generations to cause a significant level of mean population phenotype-environment match. Importantly, with matching habitat choice, the preferential sorting across the environment also leads to substantial inter-individual variability in the distance and direction of dispersal, which creates specifically directed nonrandom gene flow in the population (Bolnick and Otto, 2013; Edelaar and Bolnick, 2012, 2019; Endler, 1977; Holt, 1987)

With these phenomena in mind, researchers have used verbal models and agent-based simulations to explore the implications of matching habitat choice for ecological and evolutionary dynamics. The most thoroughly developed part of the resulting theory concerns the partitioning of genetic variance across space and its consequences. Indeed, theory suggests that non-random (directed) gene flow induced by matching habitat choice can compensate for, or even reverse, the homogenizing (maladaptive) effects of gene flow caused by diffusive (random) dispersal. This effect can reduce standing genetic variation within locally adapted subpopulations, while increasing genetic divergence among subpopulations (Bolnick and Otto, 2013; Cote et al., 2017; Edelaar and Bolnick, 2012, 2019; Edelaar et al., 2008; Felsenstein, 1976; Garant et al., 2005; Hedrick, 1986; Holt, 1987; Jacob et al., 2015, 2017; Kawecki and Ebert, 2004). At the metapopulation scale, the enhanced genetic differentiation in turn can indirectly cause assortative mating, by decreasing reproductive interactions among local populations. The resulting reproductive isolation and phenotypic segregation can eventually lead to sympatric speciation and increased biodiversity (Berner and Thibert-Plante, 2015; Bolnick and Otto, 2013; Coyne, 1992; Edelaar et al., 2008, 2023; Endler, 1977; Holt, 1987; Kirkpatrick and Ravigné, 2002; Maynard Smith, 1966; Nicolaus and Edelaar, 2018; Ravigné et al., 2009; Rice and Hostert, 1993; Rice and Salt, 1988, 1990). In addition, the adaptation promoted by phenotype-environment matching can enhance species persistence under temporal climatic changes in the environment (Bonte et al., 2012; Edelaar and Bolnick, 2012, 2019; Jacob et al., 2017; Nicolaus and Edelaar, 2018; Pellerin et al., 2019).

By contrast, predicting how matching habitat choice will influence the range dynamics of a species in a heterogeneous environment is difficult. This is partly because under matching habitat choice, increased local competition between phenotypically similar individuals counterbalances the adaptive effects of directed gene flow on fitness. In spatially structured populations, such interactions can be essentially different in central versus marginal populations. Non-quantitative predictions of the consequences of matching habitat choice on range dynamics can thus fail or mislead—even at the basic level of indicating whether its effects are substantial enough to warrant attention. Rather, quantitative theory is needed for understanding the net effect of matching habitat choice on adaptive evolution. For predicting such outcomes as the speed and ultimate extent of range shifts, developing the theory is crucial.

### 1.3 Present Work: When and How Matching Habitat Choice Influences Species Range Evolution

In the present work we propose and numerically study a deterministic partial differential equation (PDE) model for the spatiotemporal dynamics of the moments of a fitness-related quantitative trait in the presence of matching habitat choice. The predictive benefit of our model is that it allows us to quantify the extent to which matching habitat choice facilitates adaptation and speeds up range expansion. We model dispersal as a combination of diffusion (random dispersal) and a particular form of matching habitat choice that we call *phenotype-optimal dispersal*. In the most idealized formalism of matching habitat choice, individuals are able to assess the entire available habitat and move to the globally optimum location. We make the still idealized, but more realistic, assumption that individuals can only locally assess their immediate surrounding environment and direct their movement toward their maximally matching neighborhood. Continuously repeating such assessments and movements, the individuals eventually settle in a location which is the best compared with other locations within their perceivable surroundings. We model this phenotype-optimal dispersal by allowing the individuals to follow the direction of a gradient in environmental optimum phenotype. Individuals who perceive both a phenotype-environment mismatch and a non-zero environmental gradient move in the direction of the gradient, which gives a maximal (local) decrease in their perceived mismatch. If the environmental gradient is zero, or the individuals are not sensitive to it, then there will be no directed dispersal in our model as all surrounding areas appear to be equally preferable to the individuals.

We aim to characterize conditions under which phenotype-optimal dispersal is or is not sufficiently beneficial to evolve in nature. In order to do so, we use our model to describe the effects of phenotype-optimal dispersal on population dynamics measures such as expansion wave speed and equilibrium population density, as well as evolutionary processes such as rates of local adaptation and phenotypic differentiation. We focus on a continuous environment that varies in space but not in time. We also explore scenarios with repeated abrupt environmental shifts or fragmented habitat. To isolate the effects of matching habitat choice, we compare the outcomes of numerical simulations including it with the outcomes of simulations that include only diffusive dispersal.

We find that in shallow to moderate environmental gradients, phenotype-optimal dispersal has only mild effects on invasion speeds and adaptation rates. By contrast, it has several noteworthy effects in strong to extreme environmental gradients. There, it substantially increases the rate and extent of local adaptation, especially at range margins. It also reduces local phenotypic variance and increases the speed of range expansions. However, it has little effect on equilibrium population densities even in the case of a strong environmental gradient. Finally, we find that matching habitat choice in this case can greatly enhance the range expansion capacity and improve the survival prospects of a population under periodic environmental fluctuations.

## 2. Model Description

We base our model on our previous work (Shirani and Miller, 2022), which was an extension of the seminal works of Pease et al. (1989), Kirkpatrick and Barton (1997), and Case and Taper (2000) and allowed us to study the adaptive range dynamics of a community of interacting species under an environmental gradient. Since in the present work we investigate the range evolution of a solitary species, we use a reduction of our previous multi-species model to a single species case. We incorporate into the model new components that represent the effects of phenotype-optimal dispersal. We describe only those details of our previous work that are necessary for a clear description of the new model. We refer the reader to our previous work (Shirani and Miller, 2022) for further details.

Before specifying our model equations, it is worth recognizing that they are rather complex. This is because they include the contributions of three eco-evolutionary processes not included in the seminal work of Kirkpatrick and Barton (1997). The first of these is, of course, phenotype-optimal dispersal itself. The second is evolving trait variance. We include this because matching habitat choice is known to have substantial impacts on trait variance, which our work is aimed to quantify. Moreover, it is known that including the evolution of trait variance yields range dynamics very different from what occurs when variance is (arbitrarily) held fixed as in (Kirkpatrick and Barton, 1997); see for example (Barton, 2001; Shirani and Miller, 2022). The third is phenotype-dependent competition. We include this because competition is widely known as a determinant of range limits. We specifically make competition phenotype-dependent so that we can quantitatively resolve possibly conflicting effects of dispersal and competition when both depend on phenotype.

We model the m-dimensional habitat of the species by an open rectangle Ω ⊂ ℝ^m^, m ∈ {1, 2, 3}. Although in general the habitat can be three-dimensional, in this study we only consider one- and two-dimensional habitats. We model the adaptive range dynamics of the species at each location *x* = (*x*_1_, …, *x*_m_) ∈ Ω and time *t* ∈ [0, *T*], where *x*_*i*_, *i* ∈ {1, …, m}, denotes the coordinate of the location *x* along the *i*th dimension of the geographic space, and *T >* 0 denotes the evolution time horizon. For this, we derive equations that govern the joint evolution of three population quantities: *n*(*x, t*) denoting the population density of the species, *q*(*x, t*) denoting the mean value of a fitness-related quantitative phenotypic trait within the population, and *v*(*x, t*) denoting the intraspecific variance of the trait.

The derivation of the equations of the model relies on a basic equation that specifies, over a small interval of time, the variation in population density of individuals with a quantitative phenotypic trait value of *p*. To present this equation, we first denote by *φ*(*x, t, p*) the relative frequency of phenotype value *p* ∈ ℝ among all individuals of the species’ population. In addition, we denote by *g*(*x, t, p*) the intrinsic^1^ growth rate of the population of individuals with phenotype value *p*. This growth rate, as given below by (5), includes a Lotka-Volterra model of intraspecific competition. We denote by *α*(*p, p*′) the competition kernel that captures the strength of per capita effects of individuals with phenotype *p*′ on the frequency of individuals with phenotype *p*. Finally, we denote by 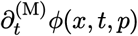 the rate of mutational changes in the frequency of phenotype *p*. The basic equation underlying the derivation of the model can then be presented as

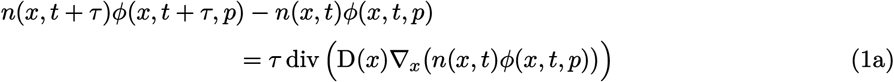

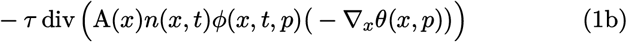

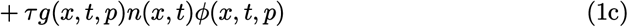

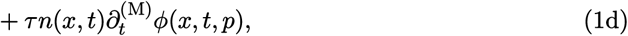

where ∇_*x*_ denotes the gradient with respect to *x*, and div denotes the divergence with respect to *x*. In writing (1), we assume that the change in the population density of individuals with phenotype *p* over a small time interval of *τ* → 0 results from the contributions of four factors: diffusive (random) dispersal of individuals to and from neighboring locations, modeled by (1a); directed (optimal) dispersal of individuals in the direction that gives them maximum environmental match, modeled by (1b); intrinsic population growth, modeled by (1c); and mutational changes in the relative frequency of *p*, modeled by (1d). As defined in Table 1, parameter D denotes the diffusion coefficient of the species’ random dispersal. Parameter A and the term −∇_*x*_*θ* in (1b) can be interpreted, respectively, as individuals’ propensity and individuals’ perceived force to disperse optimally. Further descriptions on these terms are provided in Section 2.3 below.

**Table 1:** Definition and plausible range of values of the parameters of the model (10)–(12). Except for A(*x*), the range of parameter values and their choice of units are the same as those proposed by Shirani and Miller (2022). Unless otherwise stated, the typical values given here are the values used in the numerical studies of Section 4.

| Parameter | Definition | Range | Typical | Unit |
| --- | --- | --- | --- | --- |
| $m$ | Spatial dimension of the geographic space | $\{1, 2, 3\}$ | 1, 2 | — |
| $D(x)$ | Diffusion coefficient of the species’ random dispersal | $[0, 25]^*$ | $1^*$ | $X^2/T$ |
| $A(x)$ | Measure of the species’ propensity to disperse optimally | $[0, 10]$ | 4 | $X^2/T$ |
| $K(x)$ | Carrying capacity of the environment<sup>†</sup> | $(0, 10]$ | 1 | $N/X^m$ |
| $R(x)$ | Maximum population growth rate of the species | $[0.1, 10]$ | 2 | $1/T$ |
| $V$ | Variance of the species’ phenotype utilization distribution | $[0.25, 25]$ | 4 | $Q^2$ |
| $S$ | Measure of the strength of stabilizing selection | $[0, 2]$ | 0.2 | $Q^{-2}/T$ |
| $U$ | Rate of increase in trait variance due to mutation | $[0, 0.2]$ | 0.02 | $Q^2/T$ |
| $Q(x)$ | Optimal trait value for the environment | $[0, \infty)$ | Linear<sup>‡</sup> | $Q$ |
| $\ \nabla_x Q(x)\ _{\mathbb{R}^m}$ | Magnitude of the gradient of the optimal trait | $[0, 10]$ | 0.2 | $Q/X$ |

\*When $m > 1$ , the range of values specified for $D(x)$ can be considered for each entry of $D(x) \subset \mathbb{R}^{m \times m}$ . Typically, $D$ is assumed to be diagonal.
<sup>†</sup>The carrying capacity $K$ is defined as the maximum population density when individuals are strongly competitive (generalist), that is, $V \gg 1$ . When $V$ is not large, the resulting competitive release lets the maximum population density exceed the value of $K$ specified here.
<sup>‡</sup>The typical value “Linear” specified for $Q$ means that $Q$ is typically considered to be a linear function of $x$ over $\Omega$ .

Below, we first provide a list of key assumptions that we have made in writing (1) and some other equations in the rest of the paper. We then give the formulation of the species’ intrinsic growth rate *g*(*x, t, p*) in (1c), and describe in detail how we model the optimal dispersal in (1b). The diffusion term (1a) that models species’ random dispersal is standard, and the formulation for the effect of mutational changes in (1d), based on Assumption (viii) below, is given in Section A.3 of our previous work (Shirani and Miller, 2022).

### 2.1 Model Assumptions

To derive the equations of our model, we make the following major assumptions on the populations’ dispersal and reproduction, as well as the elements of the intrinsic growth rate and the intraspecific competition kernel:

i. Random dispersal of the individuals in the habitat is diffusive.
ii. An individual’s environmental *potential energy* for optimal dispersal is proportional to the square of the difference between its phenotype value and the environment’s optimum phenotype. This assumption is crucial for the quantitative outputs of the model, as detailed in Section 2.3.
iii. Nonlinear environmental selection for an optimal phenotype Q is stabilizing for all *x* ∈ Ω. The optimal phenotype can vary over space and time.
iv. The frequency of phenotype values within the species is normally distributed at each occupied point in space at all times. That is,

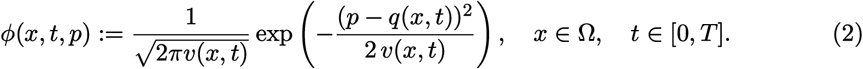
v. In the absence of selection, the intrinsic growth rate of the individuals with phenotype *p* follows a logistic growth (see (5a)) with phenotype-dependent competition. The carrying capacity and maximum growth rate of the individuals are independent of their phenotype.
vi. Environmental resources vary continuously along a resource axis. After identifying the resource axis with a phenotype axis, as described in Remark 2.2 below, the *phenotype utilization distribution* for an individual with phenotype *p* is assumed to be normal, given by

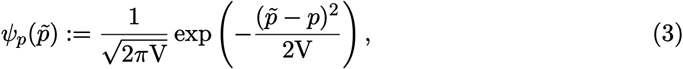

where V denotes the variance of phenotype utilization by every individuals of the species.
vii. The strength of intraspecific competition between individuals is determined by the overlap between their phenotype utilization curves, which results in the competition kernel

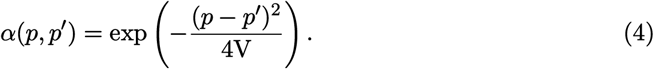 The details of the derivation of this kernel function are available in Sections A.2 and A.4 of our previous work (Shirani and Miller, 2022). Here, we only consider symmetric competition between the individuals of different phenotypes, that is, *κ* = 0 in the formulation given in our previous work.
viii. The probability of mutational changes from one phenotype *p* to another phenotype *p*′ depends on the difference *δp* = *p* − *p*′ between the phenotypes. Letting *ν*(*δp*) denote the probability density of such mutational changes, we further assume that *ν* follows a distribution with constant zero mean and constant variance V_m_. See Section A.3 of our previous work (Shirani and Miller, 2022) for the formulation of our model of mutational changes.

#### Remark 2.1 (Reproduction types)

For an isolated population, in the absence of dispersal, competition, selection, and mutations, the phenotype-density evolution equation (1) with the intrinsic growth rate (5) gives *∂*_*t*_ *n*(*x, t*)*φ*(*x, t, p*) = R(*x*) *n*(*x, t*)*φ*(*x, t, p*), where *∂*_*t*_ denotes the partial derivative with respect to *t*. This means that, under our assumption that the maximum growth rate R is phenotype-independent (Assumption (v)), the intrinsic rate of change in the density of an isolated population of individuals with phenotype *p* depends only on the population density of individuals with the same phenotype *p*. Although this condition fits in trivially with an asexual reproduction system, it also holds approximately when reproduction is predominantly by assortative mating. We note that, under our assumption of normal (symmetric and uni-modal) phenotype distribution (Assumption (iv)), a sufficiently panmictic population also does not substantially violate the reproduction assumption of our model.

#### Remark 2.2 (Phenotype Utilization Distribution)

The phenotype utilization distribution function 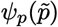 can be interpreted as a function that gives the probability density that an individual with phenotype *p* will utilize a unit of resource that is expected to be mostly utilized by (is most favorable for) an individual with phenotype 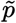. To see this more precisely, we assume (as in Assumption (vi)) that environmental resources vary continuously along a resource axis parameterized by a variable *r*. Let *ψ*_*p*_ be a probability density function that denotes the *resource utilization distribution* of individuals with phenotype *p*. That is, *ψ*_*p*_(*r*) gives the probability density that an individual with phenotype *p* obtain a unit of resource from a point *r* on the resource axis. It is then convenient for our modeling purposes, and for trait-based niche conceptualizations (Ackerly and Cornwell, 2007; Violle and Jiang, 2009), to assume that the resource axis can be identified by the phenotype axis (Roughgarden, 1979: Equ. (24.51)). This means that, we assume there exits a smooth one-to-one map *I* : *p* 1→ *r*_*p*_ that identifies the *r*-axis with the *p*-axis. That is, for every *r* ∈ R, there exist a unique phenotype 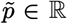 such that 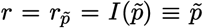. For instance, 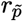 can be the point on the *r*-axis from which individuals of phenotype 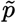 obtain their average amount of resources. Following this identification, the resource utilization distributions *ψ*_*p*_(*r*) are translated into phenotype utilization distributions 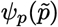, which we assume to take the Gaussian form (3) with utilization variance V. This resource-phenotype identification can be conceptualized to represent the empirical relationship between functional response traits and environments—inspiring our choice of optimal dispersal potential energy function (6) described below. In a trait-based niche quantification framework (Ackerly and Cornwell, 2007; Violle and Jiang, 2009) the variance of phenotype utilization distributions can be used to quantify within-phenotype component of a species’ niche breadth. Further details can be found in Appendix A.2 and Section 3.2 of our previous work (Shirani and Miller, 2022).

### 2.2 Intrinsic Growth Rate

In the absence of dispersal and genetic mutations, the local population dynamics of the species is determined by its intrinsic population growth, which we model as (Shirani and Miller, 2022: Equ. (17)),

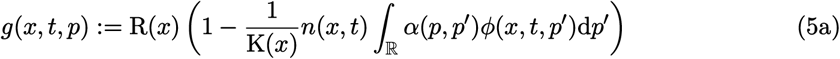

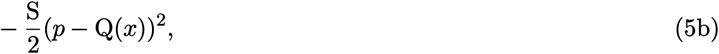

where R denotes the maximum growth rate of the species, K denotes the carrying capacity of the environment, S denotes the strength of stabilizing selection, and Q denotes the environment’s optimal trait value. The phenotype distribution *φ* and the competition kernel *α* are given by (2) and (4), respectively. The convolution term in the Lotka-Volterra model (5a) captures the effect of intraspecific phenotypic competition on the frequency of phenotype *p*. The quadratic term (5b) incorporates the effects of directional and stabilizing selection on individuals with phenotype *p*, by penalizing the phenotypes that are far from the optimal phenotype Q(*x*).

### 2.3 Phenotype-Optimal Dispersal

We model the species’ phenotype- and environment-dependent optimal dispersal by the advection term (1b). The parameter A is analogous to the *mobility* parameter that is often used in drift-diffusion models of particles flowing in a fluid. In our model, we can reasonably interpret A as a simplified model of individuals’ propensity to disperse optimally. The evolution of a species’ dispersal propensity depends on many factors, such as costs and benefits of different stages of dispersal, environmental conditions, and frequency-dependent eco-evolutionary processes (Bonte et al., 2012; Clobert et al., 2009). Moreover, the dispersal propensity of the individuals of a species is often plastic, and can change on an ecological timescale. We do not include the evolution of the species’s optimal dispersal propensity in our model though, since quantitative estimates of dispersal propensity that allow for a sufficiently meaningful approximation of its evolutionary dynamics are currently lacking in the literature. Instead, we perform our studies with different values of A to explore whether different degrees of dispersal propensity will be sufficiently beneficial to evolve in a species. Although the equations of our model allow for A to be dependent on space, in the results we present in this work we assume A to be constant in space and time.

The term −∇_*x*_*θ*(*x, p*) in (1b) is analogous to the force acting on particles in drift-diffusion models, derived from an external potential energy *θ*(*x, p*). In our model, *θ*(*x, p*) is interpreted as individuals’ perceived *dispersal potential energy* (for directed dispersal), which depends on both the phenotype *p* of each individual and the (perceived) environmental trait optimum. As a result, the advective optimal dispersal (1b) in our model represents an informed dispersal strategy that is both phenotype-dependent (dependence on *p*) and condition-dependent (dependence on Q(*x*)), as defined by Clobert et al. (2009). In this informed dispersal context, the phenotype value of an individual is an internal state of the individual, developed by the individual’s self perception. The environmental trait optimum and its gradient, as described below, are external factors that can initiate and direct the species’ optimal dispersal through the evolution of sensory and cognitive processing mechanisms in the individuals.

To fix ideas, we consider the following simplified yet meaningful model for the dispersal potential energy function of an individual with phenotype *p* at habitat location *x*,

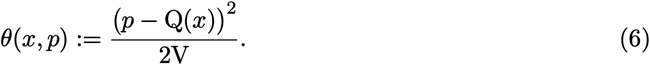

Note that this potential energy function can be obtained as a first-order approximation^2^ of the function 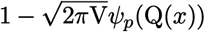, where *ψ*_*p*_ is the phenotype utilization distribution (3). That is,

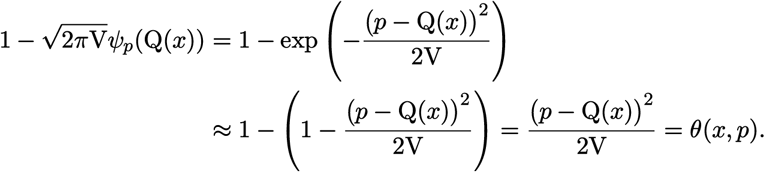

This approximation implies that the dispersal potential energy for an individual with phenotype *p* can be interpreted as how well or poorly the individual can utilize the environment’s optimal phenotype^3^ at its current location. If the individual’s phenotype matches the optimum phenotype perfectly, then there is no phenotypically and environmentally induced force on the individual to disperse. If the individual’s phenotype differs significantly from the environment’s optimum—measured relative to the species’ phenotype utilization variance V—then the individual perceives a high potential energy to disperse to habitat locations of better quality that match its phenotype. Yet, a high dispersal potential energy *θ*(*x, p*) does not generate a significant dispersal force −∇_*x*_*θ*(*x, p*) on the individual, unless a sufficiently large gradient in the dispersal potential energy is perceived by the individual. As the following discussion shows, such a gradient is present if the environmental gradient ∇_*x*_Q is sufficiently steep and the individual is sufficiently sensitive to it. In this case, the individual enjoys a considerably better-matching habitat after carrying out the directed dispersal.

The dispersal potential energy (6) gives the *optimal dispersal force* term −∇_*x*_*θ* used in (1b) as

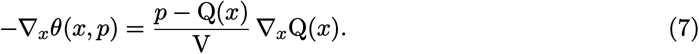

If the magnitude of the environmental gradient 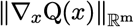 is zero (where 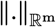 denotes the Euclidean norm in ℝ^m^) or is perceived as equal to zero due to insensitivity of an individual to the gradient, then the perceived directed dispersal force on the individual is zero—regardless of the presence of a phenotype-environment mismatch *p* ≠ Q(*x*). In this case the individual only disperses randomly, due to the diffusion term (1a). If 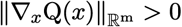, then an individual whose phenotype *p* does not perfectly match the optimum phenotype will perceive a force “pushing” it to disperse optimally. If *p >* Q(*x*), the optimal dispersal will be in the direction of the environmental gradient so that the individual disperses to habitat locations with larger optimum phenotype values, which give the individual a better phenotype-environment match. Similarly, if *p* < Q(*x*), the optimal match will be achieved by the individual through dispersal in the opposite direction of the environmental gradient. Although not possessing a true interpretation as a potential—unlike the dispersal potential energy *θ*(*x, p*) given by (6), which indeed induces a conservative force −∇_*x*_*θ* in the advection term (1b)—for ease of reference we refer to the term (*p* − Q/V) in (7) as the *phenotypic potential* for optimal dispersal.

A biological organism may not develop a perception of the dispersal force (7) in its exact mathematical sense. The phenotypic potential (*p* − Q)/V and the environmental gradient ∇_*x*_Q are perceived only approximately, and to the level needed for individuals’ decision and plan for dispersal. However, to make the derivation of the equations of our model feasible, we assume that the perceived valued of the phenotypic potential is equal to its exact mathematical value (*p* − Q)/V; see Remark 2.4 below. Yet, our derivations allow for a more reasonable perception of the environmental gradient, which we denote by 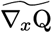 and describe in Remark 2.3 below.

Replacing the actual environmental gradient in (7) with its perceived value, we then write the individual’s *perceived dispersal force* for phenotype-optimal dispersal as:

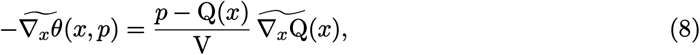

which is the dispersal force we substitute into (1b) to derive the equations of our model ((10)–(12)) given in Section 2.4.

#### Remark 2.3 (Perceived environmental gradient)

The magnitude of the dispersal force perceived by an individual is unlikely to remain directly proportional to the magnitude of the environmental gradient 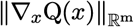 when the gradient becomes increasingly steep. When there exists a phenotype-environment mismatch *p* ≠ Q, a sufficiently steep environmental gradient should be enough to generate a maximal force on the individual to disperse. Developing proportional sensitivity to steeper gradients would then be unnecessarily costly for the individual, as it would not change the individual’s dispersal behavior any further. To approximately incorporate such information saturation in individual’s perception of the environmental gradient, we replace ∇_*x*_Q in (7) by the following *perceived gradient*:

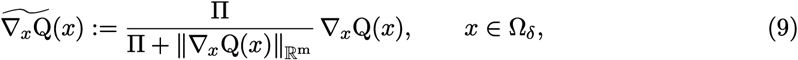

where Π is a constant that denotes the maximum perceived magnitude of the environmental grtadient. When 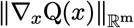 is much smaller than Π, the perceived gradient approximately equals the actual gradient. When 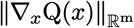 is much larger than Π, the magnitude of the perceived gradient approximately saturates to the maximum value Π, but its direction will always be the same as the direction of the actual gradient. The smaller habitat Ω_*δ*_ specified in (9) includes all points of Ω except those that are closer than a constant *δ* to the boundary of Ω. This is to avoid the complexity of exposition here (due to the indefiniteness of the gradient at boundary points,) and to allow for simpler boundary conditions. In Appendix A, we provide details on how to smoothly extend 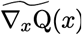 in (9) to the whole habitat Ω, based on the assumption that individuals can sense the habitat boundary and avoid crossing it. Specifically, for the one- and two-dimensional habitats that we simulate in this work, we use the definitions (21) and (22) given in Appendix A for 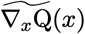 over the entire habitat Ω. Denoting the units of space and trait by X and Q, respectively, we assume in all numerical simulations that the individuals can perceive the habitat boundary when they get as close as *δ* = 2 X to the boundary. Moreover, we set the maximum perceived gradient to be Π = 1 Q/X, which still corresponds to a relatively steep environmental gradient, as we discussed in Section 3.2 of our previous work (Shirani and Miller, 2022). In our simulations, we use the dispersal propensity parameter A as an adjustment parameter for the species’ total rate of optimal dispersal.

We note that the perceived gradient 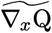 can also be interpreted, in some sense, as the sensitivity of the individuals’ dispersal force to changes in habitat quality. In nature, individuals of a species may develop a perception of the environmental gradient not only by directly sensing their environment’s conditions, but also by collecting information through exploration (Armsworth and Roughgarden, 2005; Selonen and Hanski, 2006) and collective behavior of their conspecifics. The diffusive movement term (1a) that we have included in our model can additionally capture such exploratory movements of the individuals for the purpose of gaining a better perception of their environment’s gradient. Tracking the environmental gradient can also occur through taxis, that is, in response to environmental stimuli (guides) that are strongly correlated with gradient of the optimum trait. For instance, changes in temperature, chemical concentration, topography, wind strength and direction, water flow, or light intensity can effectively canalize the movement of individuals in the direction of the environmental gradient (Baguette et al., 2013).

#### Remark 2.4 (Perceived phenotypic potential)

An individual with the ability to locate and disperse to a matching habitat most likely can develop a reasonably accurate self-assessment of the mismatch between its phenotype and the environmental optimum, *p* − Q(*x*), or equivalently, a perception of the phenotypic potential (*p* − Q(*x*))/V at every location *x* in the habitat. We can model the perceived value fo the phenotypic potential as a function *f* ((*p* − Q)/V). For example, *f* could reasonably be a saturation function similar to what we considered for the magnitude of the environmental gradient in Remark 2.3. However, the presence of such a nonlinear function of *p* in our model of optimal dispersal makes the derivation of the final equations of the model infeasible, or unnecessarily complicated. An alternative choice without over-complicating the derivations would be in the form 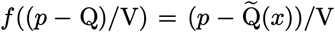, where 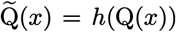 gives the perceived value of the trait optimum as a function *h* of its actual value. This particularly means that we assume the individual’s assessment of its own phenotype is exact. Substituting this perception of the phenotypic potential for the exact value (*p* − Q(*x*))/V in (8) would simply result in replacing Q(*x*) with its perception 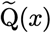 in the optimal dispersal terms (10b), (11e), (12e) of the model equations (10)–(12) given in Section 2.4. However, since we assume the perception of *p* is exact and it is likely that the ability to perceive the environmental optimum Q evolves jointly with the self-perception of *p*, we choose to set 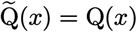 in all of our equations. This means that throughout this work we always assume that the perception of the phenotypic potential is exact.

### 2.4 Model Equations

The basic equation (1), and its components described above, provide the ingredients that we need for deriving the equations of our model for the joint evolution of a species’s population density *n*(*x, t*) and the mean value *q*(*x, t*) and variance *v*(*x, t*) of a quantitative fitness-related trait within the species’ population. Assumption (iv) is a key assumption in developing our model as it allows for an exact moment closure in deriving the equations of trait mean and trait variance. In this section, we only present the final equations of our model. The derivation of the equations is provided in detail in Appendix B.

Before presenting the equations of the model, we note that the definitions of all model parameters and their plausible ranges of values are given in Table 1. Note that D(*x*) ∈ ℝ^m*×*m^, whereas the rest of the parameters are scalar-valued. Also, S, U, and V are assumed to be constant throughout the habitat, whereas D, A, K, R, and Q can be variable in space. Although their dependence on *t* is not explicitly shown in the equations, all these model parameters can also vary in time. A discussion of the choice of parameter units and their plausible values is provided in Section 2.5 below.

Now, letting *∂*_*t*_ denote the partial derivative with respect to *t*, our equation for the evolution of the species’ population density *n*(*x, t*) for all *x* ∈ Ω and *t* ∈ [0, *T*] is given as

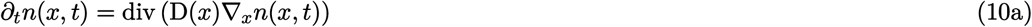

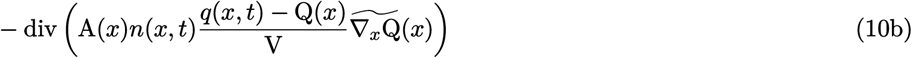

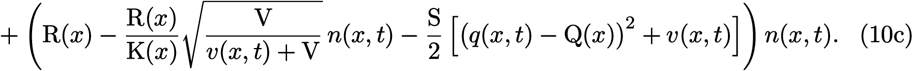

Further, letting 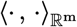 denote the standard inner product in ℝ^m^, our equations for the population’s trait mean *q*(*x, t*) and trait variance *v*(*x, t*) are

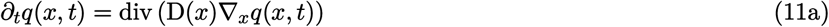

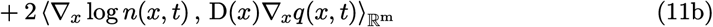

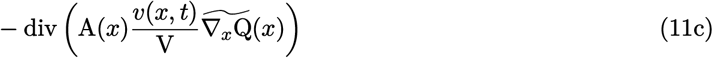

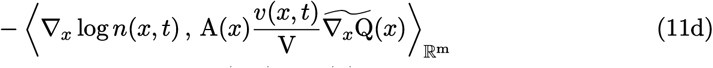

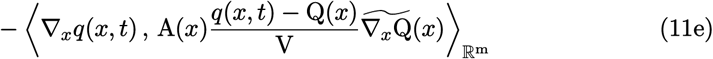

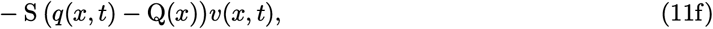

and

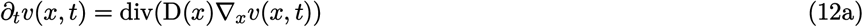

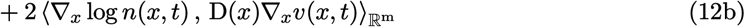

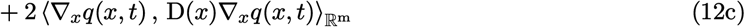

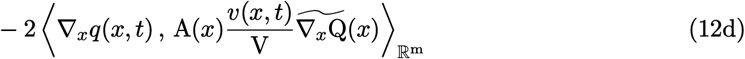

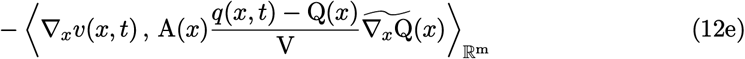

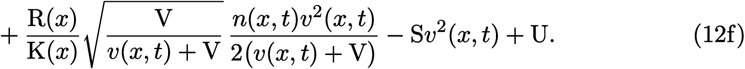

For our simulations of a one-dimensional habitat Ω = (*a, b*), we assume no phenotype flux through the habitat boundary. Since the perceived environmental gradient 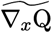 given by (21) in Appendix A has no component normal to the boundary, the advection term (1b) does not result in any phenotype flux through the boundary. Therefore, our no-flux boundary assumption simply implies the homogeneous Neumann (reflecting) boundary conditions

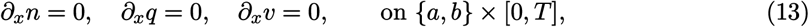

which is the boundary condition we discussed in Remark 1 and Appendix A.5 of our previous work (Shirani and Miller, 2022). For the two-dimensional habitat Ω = (*a*_1_, *b*_1_) × (*a*_2_, *b*_2_) that we simulate in this work, we set reflecting boundary condition at boundary lines *x*_1_ = *a*_1_ and *x*_1_ = *b*_1_, and we assume that the habitat is extended periodically in the *x*_2_-direction.

In comparison with the equations of the single-species model that we studied in Section 4 of our previous work, the inclusion of the optimal dispersal strategy in the present work results in the additional term (10b) in the equation for population density, the terms (11c)–(11e) in the equation for trait mean, and the terms (12d) and (12e) in the equation for trait variance. As a result, all quantitative measures of population dynamics, such as population density, speed of range expansion, local adaptation rates at both the range center (core) and range margins, asymmetric core-to-edge gene flow, and the dynamics of intraspecific trait variance are expected to be affected by individuals’ ability to disperse optimally to matching habitats. We demonstrate such impacts under different evolutionary regimes in our computational studies presented in Section 4.

### 2.5 Units and Parameter Values

We use the same units as we discussed in our previous work for the quantities included in our model. Specifically, we denote the unit of time by T and we set 1 T to be equal to the mean generation time of the species. For a one-dimensional habitat, we choose the unit of space so that the diffusion coefficient D of the population becomes unity. That is, denoting the unit of space by X, we set 1 X to be the root mean square of the (random) dispersal distance of the population in 1 T, divided by 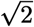. Estimates of the random component of the species’ dispersal can be obtained, for example, by measuring dispersal distance of a subpopulation of individuals that are well adapted to the environment at the core of the population. Due to their negligible phenotype-environment mismatch, such individuals do not perceive a significant force compelling directed dispersal. For multi-dimensional habitats, the same approach can be used to set the unit of space for each spatial dimension independently. Moreover, we denote the unit of measurement for population abundances by N. Having set the unit of space, we set 1 N to be equal to the carrying capacity of the environment for 1 X^m^ unit of habitat volume. This results in the carrying capacity becoming equal to unity. Finally, we denote the unit of measurement for the quantitative trait by Q, and we set 1 Q to be equal to one standard deviation of the trait values at the core of the population. Further discussion of our choices of units is available in Section 3.1 of our previous work (Shirani and Miller, 2022).

Our suggestions of plausible ranges of values given in Table 1 are discussed in detail in Section 3.2 of our previous work, except for the new parameter A. The values specified as “typical” in Table 1 are the values we use as default parameter values in our numerical simulations when not otherwise stated. Due to the level of abstraction that is inevitably present in our model of optimal dispersal, finding a biologically reasonable range of values for the dispersal propensity parameter A based on empirical measurements available in the literature is infeasible. Instead, we investigate a range from 0 to 10 X^2^/T by simulating the model with different values of A and observing the resulting range of variation in the population density, speed of range expansion waves, and magnitude of the trait variance. In an extreme situation, for example, if the mean value of the trait in (10b) differs from Q by one phenotype utilization variance and the magnitude of the environmental gradient is sufficiently greater than the preset value Π = 1 Q/X, then a maximum dispersal (advection) rate of approximately 10 X/T is created in a one-dimensional habitat at propensity value A = 10 X^2^/T.

## 3. Interpretation of the Model Equations

Before presenting our numerical results, we discuss the interpretation of each of the terms involved in the equations. Inspecting the equations provides useful mechanistic insight into the less-intuitive impacts of phenotype-optimal dispersal.

### 3.1 Population Density Equation

The diffusion term given by (10a) in our model gives the rate of change in the population density of the species due to random dispersal. In addition to modeling exploratory movement of individuals for developing perceptions of the environment, as we discussed before, this diffusion term can also incorporate the effects of other uniformed movements, such as short-range movements to escape kin competition or inbreeding (Clobert et al., 2009). Moreover, the presence of this random dispersal term in the equations further makes it possible for local populations to leave a locally optimal location and eventually settle in a location that globally maximize their phenotype-environment match.

The advection term (10b) incorporates changes in population density due to directed optimal dispersal. When the mean phenotypic potential for optimal dispersal, (*q* −Q)/V, and the perceived environmental gradient 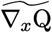 are both non-zero, the population density undergoes a directional change. If (*q* − Q) *>* 0, the whole population moves in the direction of the environmental gradient. If (*q* − Q) *<* 0, the population moves in the opposite direction. In either case, the mean mismatch |*q* − Q| is reduced through the directed movement.

The term 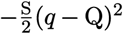 in (10c) shows the effects of natural selection in reducing the population density when the trait mean *q* differs from the optimum Q. The term 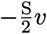 in (10c) captures the effect of the phenotypic load imposed by natural selection on the population growth, compared with a monomorphic population. Note that greater values of trait variance create stronger phenotypic loads. The term 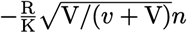 in (10c) represents the decrease in population growth rate due to the average intraspecific competition between the individuals. Unlike the phenotypic load, the average intraspecific competition becomes weaker when the trait variance *v* becomes larger. This is because larger trait variance implies greater average difference between phenotypes and hence less competition load due to (4). Furthermore, when individuals’ phenotype (resource) utilization variance V becomes smaller, that means when the individuals become more specialized, the average competition becomes weaker. This is because there is a lower chance that specialists will utilize the same resources. Note that competitive release at small values of V can allow for population density to increase significantly above K—which is the carrying capacity we define for sufficiently competitive (generalist) individuals with V ≫ 1.

### 3.2 Trait Mean Equation

The terms (11a) and (11b) show how random gene flow caused by diffusive dispersal affects the rate of change of trait mean, or equivalently, the local adaptation rate of the population. The divergence term in (11a) represents the homogenizing effect of random gene flow. Since population density changes sharply at the range margin, ∇_*x*_ log *n* in (11b) is significantly larger near the edge of the population, compared with the core. Therefore, (11b) effectively models asymmetric core-to-edge gene flow caused by random dispersal. In the absence of optimal dispersal, such maladaptive gene flow can potentially result in gene swamping at marginal populations (Kirkpatrick and Barton, 1997; Lenormand, 2002).

The terms (11c)–(11e) capture the effects of individual-level optimal dispersal on population-level local adaptation. For the divergence term (11c), we can use the product rule for the divergence of a scalar field times a vector field^4^ (with the scalar field *v*) to write

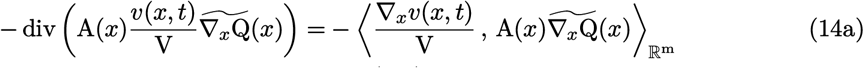

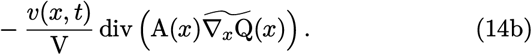

The inner product term (14a) implies that, due to directed movements, the trait mean in a local population decreases when the gradient in population’s trait variance is aligned, or makes an acute angle, with the gradient in trait optimum. The trait mean increases if the two gradients make an obtuse angle, or point in opposite directions. The simulation results provided in our previous work (Shirani and Miller, 2022: Figure 2) and the results given in Figure 1 below, as well as empirical observations (Takahashi et al., 2016), show that trait variance during the range expansion of a species—over a continuous habitat with linearly varying trait optimum—decreases from core to edge. This means that, trait variance gradient will point in the same direction as of the environmental gradient on one side of the population’s range, whereas it will point in the opposite direction on the other side. As a result, changes in trait mean due to (14a) will be increasing on one side and decreasing on the other side. Such changes are often adaptive. For instance, due to (14a), the trait mean in Figure 1 will increase in the right half of the population’s range, and will decrease in the left half. In both cases, the population gets better adapted to the environment optimum.

**Figure 1:**
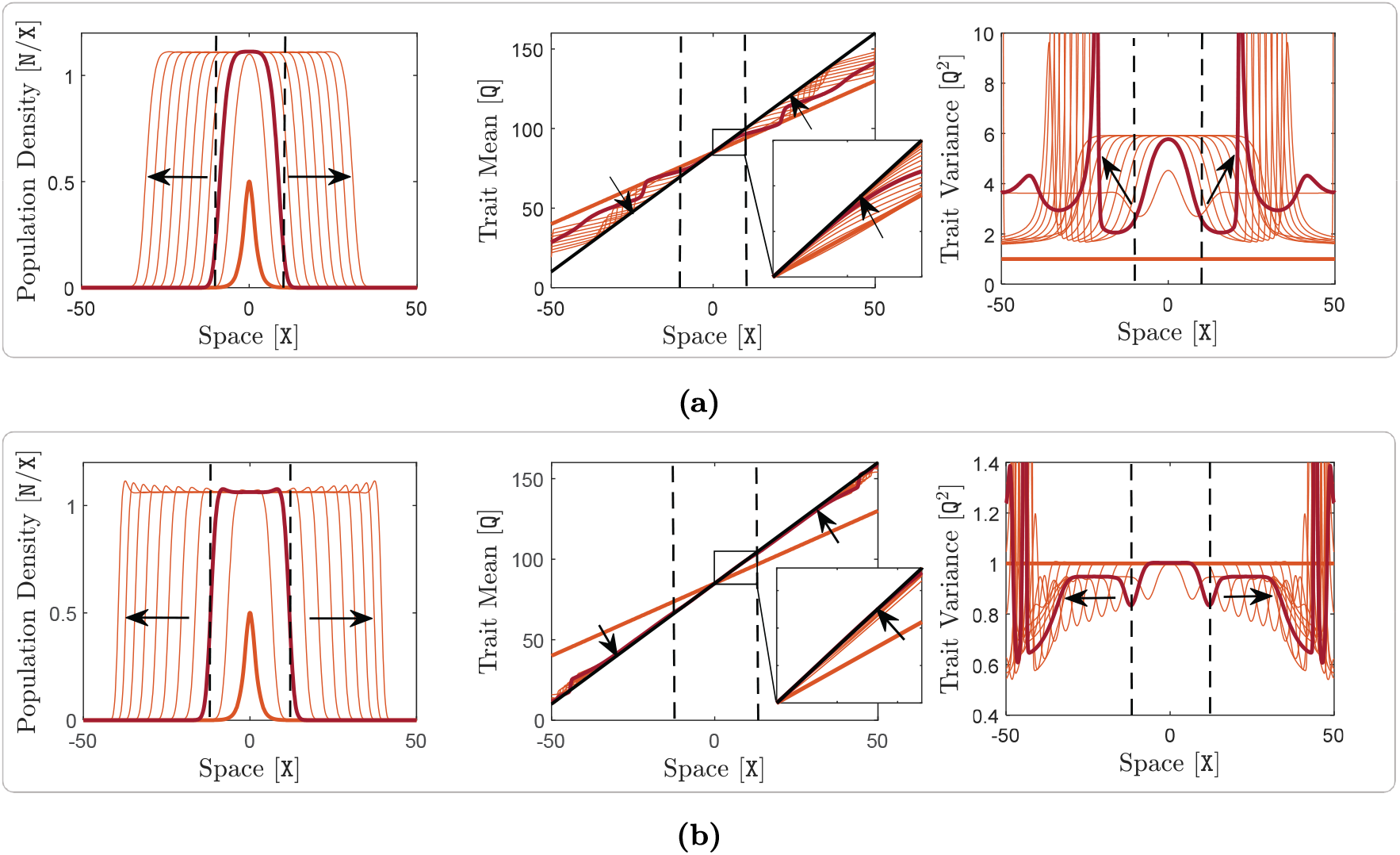
Adaptive range dynamics of a species in a one-dimensional habitat with steep environmental gradient. Here, m = 1, Q(*x*) is linear in *x* with a relatively steep gradient of ∇_*x*_Q = 1.5 Q/X, and A(*x*) is constant, taking different values in panels (a) and (b). The rest of the model parameters take the typical values given in Table 1. In each of the panels, evolution of the population density *n*(*x, t*) is shown on the left, evolution of the trait mean *q*(*x, t*) is shown in the middle, and evolution of the trait variance *v*(*x, t*) is shown on the right. Panel (a) shows the range expansion dynamics of a species without optimal dispersal, A = 0 X^2^/T, whereas panel (b) shows the range expansion dynamics of the species with strong optimal dispersal, A = 10 X^2^/T. In all graphs, curves are shown at every 4 T, and the thick orange curves indicate the initial curves at *t* = 0 T. In the insets of trait mean graphs, curves are shown at every 1 T. Arrows show the direction of evolution in time. In each graph, a sample curve at *t* = 8 T is highlighted in red. Dashed lines indicate the effective edges of the population at *t* = 8 T, associated with the inflection points on the highlighted curve of population density. The solid black lines in the graphs of trait mean show the environmental trait optimum Q. The curves of trait variance take large values outside the effective range of species. Such values are not biologically meaningful as they do not occur within the range of the species, and have been cut to smaller values for better visualization of the meaningful parts of the graphs.

**Figure 2:**
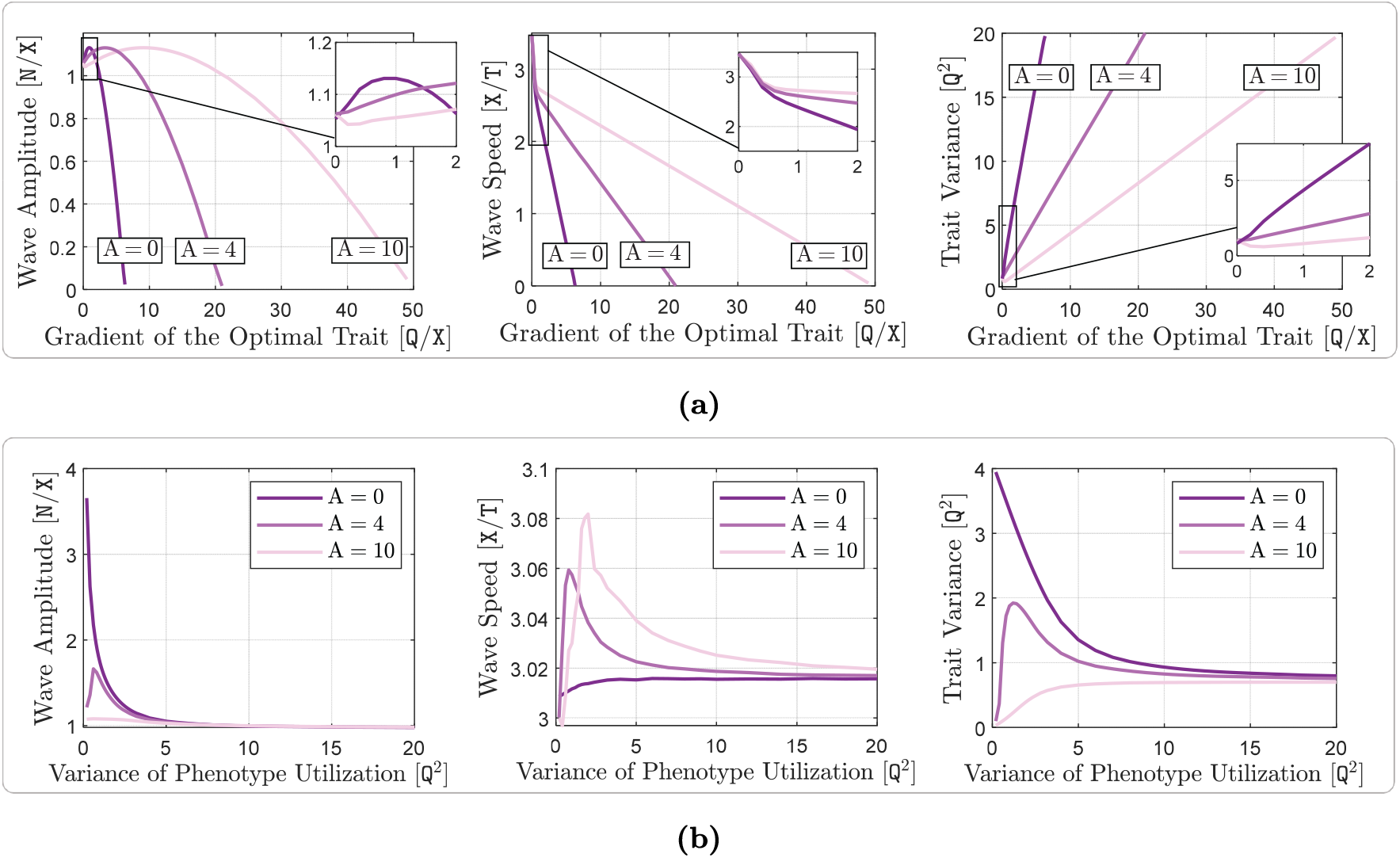
Effects of optimal dispersal on range expansion waves and maximum intraspecific trait variance of a species. Here, m = 1, and A(*x*) takes three different constant values, 0 X^2^/T, 4 X^2^/T, and 10 X^2^/T. The trait optimum Q(*x*) is linear in *x*, with variable gradient in panel (a) and constant gradient of ∇_*x*_Q = 0.2 Q/X in panel (b). The phenotype utilization variance takes the constant value V = 4 Q^2^ in panel (a), and is variable in panel (b). The rest of the model parameters take their typical values given in Table 1. In each panel, variations in the amplitude of the traveling waves are shown on the left, variations in the speed of the traveling waves of population density are shown in the middle, and variations in the maximum intraspecific trait variance are shown on the right. Panel (a) shows the effects of different levels of optimal dispersal at different magnitudes of the environmental gradient ∇_*x*_Q. Panel (b) shows the effects of different levels of optimal dispersal at different values of the individuals’ phenotype utilization variance V.

The component (14b) of (11c) shows that divergence in the perceived environmental gradient can indeed result in changes to the trait mean, even if population density and trait variance are spatially homogeneous. When A(*x*) is constant, as we assume throughout this work, trait mean is decreased by (14b) if div 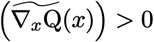. An illustration of the movement rates and directions that result in such a decrease in trait mean is provided in Figure S1. Similarly, trait mean increases when div 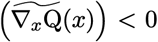. Knowing that div 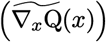 is in fact the (perceived) Laplacian of the trait optimum at *x*, we can also think of the effect of (14b) as changing the trait mean in an opposite direction to the local average change^5^ in the trait optimum. Depending on the value of the trait mean, and in particular on whether it is below the trait optimum or above it, such changes can be adaptive or maladaptive to the population. However, even if the pure effect of divergence in perceived environmental gradient appears to be maladaptive, its combined effect with the several other components of the optimal dispersal present in the model—which jointly affect population density, trait mean, and trait variance—can still be adaptive. In this work, however, we always assume that the environmental gradient is constant in both space and time. This implies that div 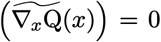 in our simulations, except in the close vicinity of the habitat boundary. Therefore, (14b) does not have any impact on the local adaptation that we observe in our results due to phenotype-optimal dispersal.

As with (11b), the presence of ∇_*x*_ log *n* in (11d) implies that this term mainly represents the asymmetric core-to-edge effects of gene flow caused by directed dispersal. To understand whether the effects of (11d) are adaptive or maladaptive, we first note that (11d) can be rewritten by taking the scalar term 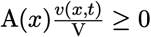 out of the inner product, giving 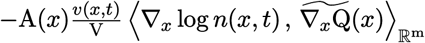. Due to the population’s adaptation to the environment during its range expansion, ∇_*x*_*q* is expected to be aligned with 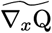, which implies that (11d) and (11b) have opposite signs. As a result, the core-to-edge directed gene flow cause by phenotype-optimal dispersal will indeed be adaptive at the range margin, unlike the maladaptive effects of the asymmetric gene flow created by random dispersal. Therefore, (11d) represents one of the main effects of matching habitat choice for facilitating adaptation at range margins. Note that, (11d) further implies that larger values of trait variance makes such adaptive effects stronger.

The inner product term (11e) shows how the mean phenotypic potential for optimal dispersal, (*q* − Q)/V, and the perceived environmental gradient directly cause local adaption. First note that (11e) can be rewritten as 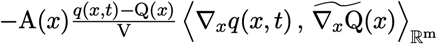. Due to the expected alignment between the directions of ∇_*x*_*q* and 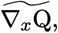, (11e) is then negative when (*q* − Q) *>* 0, and it is positive when (*q* − Q) *<* 0. In either case, the resulting change in trait mean due to (11e) is adaptive, that is, it decreases the magnitude of the mismatch |*q* −Q|. Importantly, (11e) explicitly shows that the rate of population-level adaptation is higher when the mean phenotypic potential of individuals for optimal dispersal is greater and their perceived environmental gradient is steeper. Note that, as the population gradually adapts to the environment, the mean phenotypic potential for dispersal decreases asymptotically. This means that, the rate of adaption caused by (11e) decreases as the population gradually gets better adapted to the environment. Yet (due to the terms (11c) and (11d) we discussed above) the presence of trait variation in the population provides an additional adaptation force independent of the mean mismatch |*q* − Q|, which works to enhance the overall rate of adaptation.

Finally, the reaction term (11f) represents local adaptation by natural selection. When (*q*−Q) *>* 0, this term forces a decreasing change in *q*, which results in a decrease in |*q* − Q| and enhanced local adaptation. Similarly, when (*q* − Q) *<* 0, the trait mean *q* increases due to (11f), leading to a decrease in |*q* − Q| and better adaptation. Importantly, (11f) demonstrates the crucial role of genetic variation in enabling adaptation by natural selection. The adaptation rate controlled by (11f) is directly proportional to the level of trait variance *v*. Larger values of *v* imply greater amounts of genetic variation for natural section to act upon, and hence faster adaption rates by natural selection.

### 3.3 Trait Variance Equation

The terms (12a)–(12c) represent the effects of random dispersal on rate of change of trait variance. Along with (11a), which tends to homogenize trait mean, the divergence term in (12a) tends to homogenize trait variance. Therefore, (11a) and (12a) together capture the homogenizing effects of random gene flow on population’s phenotypes. As with what we described above for the trait mean equation, the presence of the term ∇_*x*_ log *n* in (12b) implies that (12b) mainly captures the effects of asymmetric core-to-edge random gene flow on trait variance. As our simulation results shown in Figure 1 below imply, ∇_*x*_ log *n* and ∇_*x*_*v* are often aligned with each other during the range expansion of a population. Therefore, (12b) is expected to be positive, and hence it tends to increase trait variance at range margins, as the range expands. This can explain a traveling-wave profile for trait variance, as we observe in our numerical simulations. The inner product term (12c) is always positive, and it depends on the magnitude of the gradient in trait mean. Since the population typically is well-adapted at its core, ∇_*x*_*q* closely follows ∇_*x*_Q at central regions of the range. As a result, (12c) is the term which is responsible for inflating trait variance at the population’s core; a well-known effect of random gene flow over an environmental gradient (Barton, 2001; Garant et al., 2007; Lenormand, 2002). The steeper the environmental gradient, the larger the trait variance at central populations.

The terms (12d) and (12e) give the effects of the non-random gene flow created by optimal dispersal on the rate of change of trait variance. Since ∇_*x*_*q* and ∇_*x*_Q are expected to be in the same direction as the population adapts to the environment, (12d) typically takes negative values. Therefore, trait variance is reduced due to (12d). Since ∇_*x*_*q* follows ∇_*x*_Q more closely in central regions, where the population is well-adapted to the environment, and since *v* is also larger in those regions, the reduction in trait variance caused by (12d) is significantly stronger at the well-adapted core of the population. The mismatch (*q* − Q) and the gradient ∇_*x*_*v* in (12e) both take larger magnitudes near the range margins, where the population is not strongly well-adapted. As a result, (12e) mainly influences the trait variance at marginal populations. As discussed in our previous work (Shirani and Miller, 2022: Sect. 2.1), and shown in Figure 1 below, the decreasing core-to-edge profile of trait variance during range expansion over a constant environmental gradient (linearly changing Q) is directly related to the spatial profile of the trait mean. Near the peripheral regions, over which maladaptive gene flow causes *q* to fall below the optimum Q, trait variance decreases in the direction of the environmental gradient. That means, when (*q* −Q) *<* 0, we expect 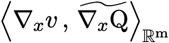 to be negative. This implies that (12e) will be negative when (*q* −Q) *<* 0. Likewise, near the peripheral regions over which (*q* −Q) *>* 0, we observe *v* to be increasing in the direction of the environmental gradient. As a result, (12e) will also be negative when (*q* − Q) *>* 0. Therefore, in any case, (12e) tends to decrease the trait variance, predominantly within less-adapted marginal populations.

The above discussion implies that the directed gene flow generated by phenotype-optimal dispersal reduces trait variance both within marginal and central populations. As we pointed out in the Introduction section, reducing phenotypic variation within local populations is in fact one of the most important predicted effects of matching habitat choice. The terms (12d) and (12e) in our model show how this reduction is controlled by the interaction between several factors, such as mean phenotypic potential for optimal dispersal (*q* −Q)/V, trait variance, perceived environmental gradient, and gradient in trait mean and variance.

The first term in (12f) models the effects of intraspecific competition on the rate of change of trait variance. Note that this term is always nonnegative, implying that competition tends to increase trait variance. This is because competition reduces the fitness of individuals with close phenotype values, while it does not significantly affect the individuals with sufficiently different phenotypes. When V → ∞, that is, when individuals become highly generalist, the completion term in (12f) vanishes to zero and causes no inflation in trait variance. This is because competition affects the fitness of highly generalist individuals almost uniformly, as these individuals almost equally utilize all available resources regardless of their phenotype. As a result, no phenotype variation is generated by the competition between them. When V → 0, that is, when individuals become highly specialist, the term 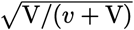 in (12f) converges to zero. However, the competitive release gained by the population when V → 0 allows for the maximum steady-state population density *n*, controlled by (10c), to take arbitrarily large values. As a result, the competition term in (12f) remains positive (non-zero) when V → 0. Our computations of spatially homogeneous steady-state values of trait variance imply that, when V → 0, trait variance increases to a finite value due to this non-zero competition term; see Figures 2b and S4b for A = 0.

The term −S*v*^2^ in in (12f) represents the well-known effect of natural selection in eroding genetic variation by eliminating less-fit individuals. Note that, the larger the trait variance *v*, the stronger the effect of natural selection, and hence the higher rate of reduction in *v*. Finally, the presence of the constant term U in (12f) shows the effect of mutation, as a perpetual source of genetic variation within the population.

## 4. Results

To determine when, if ever, and how phenotype-optimal dispersal markedly affects range dynamics and adaptation, we solve the equations of the model (10)–(12) numerically using the parameter values given in Table 1. Our discussion in Section 3 indicates how each of the eco-evolutionary forces incorporated into the model influences, separately, the species’ range evolution. However, numerical studies are necessary since intuition is inadequate for predicting how these forces interact to determine population density and trait moment dynamics. Moreover, a rigorous analytical study of our model is infeasible, due to its level of complexity.

In all except one of our numerical studies we consider a one-dimensional continuous habitat with linearly changing environmental optimum phenotype. Our only study in a two-dimensional habitat investigates the effects of phenotype-optimal dispersal in presence of habitat fragmentation. The details of the numerical scheme we use to compute the solutions are given in Appendix B of our previous work (Shirani and Miller, 2022). A discussion of the challenges in numerical computation of the solutions can also be found in Section 6.5 of our previous work.

### 4.1 Adaptive Range Dynamics with Phenotype-Optimal Dispersal

We first demonstrate how optimal dispersal changes the spatiotemporal population dynamics of a species. To isolate the effects of phenotype-optimal dispersal, we carry out pairs of numerical simulations that are identical except that only one incorporates this form of dispersal. Specifically, in one of the simulations we only consider random dispersal, which means that we set A = 0 X^2^/T. In the other simulation we additionally include strong optimal dispersal, by setting A = 10 X^2^/T. The other parameters of the model remain the same in both simulations. Other than the trait optimum Q, which is considered to be linearly increasing over Ω, the rest of the parameters are assumed to be constant. We consider a one-dimensional habitat Ω = (−50 X, 50 X) ⊂ ℝ with the reflecting boundary conditions (13).

#### 4.1.1 Range Evolution in Steep Environmental Gradients

To make the effects of optimal dispersal strong enough to be clearly visible in our graphs, we consider a steep environmental gradient of d_*x*_Q = 1.5 Q/X. We initially introduce the species at the center of the habitat with a density given as *n*(*x*, 0) = 0.5 sech 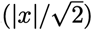. We assume that the trait mean in this initial population varies linearly in space, with the constant gradient of ∇_*x*_*q*(*x*, 0) = 0.6 ∇_*x*_Q(*x*). We further assume that the initial population is perfectly adapted to the environment at the center, *q*(0, 0) = Q(0), and has a constant trait variance of *v*(*x*, 0) = 1 Q^2^. The results of our simulations over the computation time horizon of *T* = 40 T are shown in Figure 1.

We observe that, with or without optimal dispersal, the species’ population density first grows to a maximum capacity—determined by the environment’s carrying capacity and the level of competitive release—and then expands indefinitely in the form of a traveling wave. The population’s trait mean converges to the optimum trait, due to the adaptation caused both by natural selection and by phenotype-optimal dispersal, when it exists. The population’s trait variance evolves a spatial profile that is decreasing from core to edge of the population. The maximum trait variance at the population core reaches a constant upper bound.

Without optimal dispersal, the peripheral populations near the wavefronts are maladapted and exhibit low trait variance compared with the population near the range center (Figure 1a). This maladaptation is mainly caused by asymmetric core-to-edge random gene flow, which decreases the trait mean below the optimum near the right edge and increases it above the optimum near the left edge. At the core, population density is almost uniform and hence gene flow is symmetric. This implies that random gene flow does not significantly affect the trait mean at central locations and local adaptation is maintained by natural selection. Since the environmental gradient is steep and the population is well-adapted to it at its core, random gene flow from adjacent areas generates large phenotypic variations among central individuals. Near the edges, however, the trait mean fails to follow the steep gradient in the optimum trait. As a results, trait variance decreases from core to edge in parallel with the decrease in gradient of the trait mean. Further descriptions of the profiles of trait mean and variance under random dispersal are available in Section 4.1 of our previous work (Shirani and Miller, 2022).

With optimal dispersal, both the rate and the extent of adaptation in central and peripheral populations are enhanced compared to the diffusion-only case (Figure 1b). The traveling-wave dynamics in the two cases are similar, but we note several differences. Comparing the insets in the graphs of trait mean in Figures 1a and 1b, we see that phenotype-optimal dispersal significantly increases the local adaptation rate of the population. Convergence to the environmental optimum at the population’s core occurs in almost one generation time with optimal dispersal, whereas the same level of adaptation takes more than ten generations to occur in the absence of optimal dispersal. The level of maladaptation at range margins is also substantially decreased by the directed gene flow created by phenotype-optimal dispersal. These are all consistent with the general predictions regarding the adaptive effects of matching habitat choice, such as those discussed in the Introduction section, or generated by inspecting the equations of our model. Importantly, by comparing the curves of trait variance in Figures 1a and 1b, we also observe that the phenotypic assortment resulting from preferential movements under optimal dispersal significantly reduces trait variance within the population. Moreover, the curves of population density in Figures 1a and 1b show that adaptation reinforced by optimal dispersal increases the overall range expansion speed of the population, especially at earlier stages of population establishment in the habitat. The maximum population density, however, is not notably affected by optimal dispersal at the environmental gradient we simulated in Figure 1.

#### 4.1.2 Range Evolution with Specialized Individuals

The phenotypic potential of individuals to disperse optimally (that is, (*p* −Q)/V in (8)) is stronger when their phenotype utilization variance V is smaller; meaning that the individuals have a higher degree of specialization. To see if phenotype-optimal dispersal by specialists can significantly influence the range dynamics when the environmental gradient is not very steep, we simulate our model with the relatively small value of V = 1 Q^2^ under the typical gradient of ∇_*x*_Q = 0.2 Q/X. The results are shown in supplementary Figure S2. We observe similar effects to those shown in Figure 1. That is, phenotype-optimal dispersal facilitates local adaptation of specialists on a within-generation timescale, reduces their average maladaptation at range margins, and reduces their within-population trait variance. However, unlike what we observed in Figure 1, the population’s range expansion speed is not considerably increased. Without optimal dispersal, we see in Figure S2a that the population density rises significantly above K = 1 N/X. This is due to the population’s ecological release gained by less-competitive specialists. However, phenotype-optimal dispersal limits the level of competitive release, as it counteracts the effects of smaller phenotype utilization variance (less competition) by significantly reducing phenotypic variation within the population. Phenotypically close individuals can still remain sufficiently competitive even when they are specialists and utilize fewer common resources. As a result, we see in Figure S2b that population density is still approximately bounded by the carrying capacity K = 1 N/X when the specialist individuals disperse optimally.

### 4.2 When Does Phenotype-Optimal Dispersal Influence Adaptive Range Dynamics?

The effects of phenotype-optimal dispersal are marked only when environmental gradients are very steep. The pronounced effects of optimal dispersal we observed in Figure 1 were obtained at the environmental gradient ∇_*x*_Q = 1.5 Q/X. The steepness of the environmental gradient is in fact a key factor in determining whether or not phenotype-optimal dispersal will be sufficiently consequential in a species’ range evolution. The perceived force for optimal dispersal (8) is directly proportional to the (perceived) magnitude of the environmental gradient. Moreover, the phenotypic potential (*p* −Q)/V in (8) is directly influenced by the level of phenotypic variation in the population, which in turn can be substantially inflated by random gene flow under steep environmental gradients. Estimates of realistic values for the slopes of environmental optimum gradients in nature, however, are not widely available in the literature—noting that our choices of units for ∇_*x*_Q requires joint measurements of optimal trait values, dispersal distance, and generation time. Yet, based on some available data, in our previous work we argued that a plausible range of values for 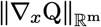 could lie approximately between 0 and 2 Q/X, (Shirani and Miller, 2022: Sect. 3.2). This means that the gradient ∇_*x*_Q = 1.5 Q/X used for the results shown in Figure 1 is very steep; for example, possibly associated with long-range dispersal of birds over elevational gradients. In supplementary Figure S3, we show our simulation results for a shallower environmental gradient of ∇_*x*_Q = 0.2 Q/X, which might be more typically observed in nature. We see that phenotype-optimal dispersal does not significantly affect the species’s range dynamics in this case.

Further simulations with varying levels of environmental gradient and individuals’ specialization confirmed that optimal dispersal was significantly consequential when gradients were very steep. In each of these simulations, we considered three different levels of optimal dispersal propensity, A = 0 X^2^/T (no optimal dispersal), A = 4 X^2^/T (medium optimal dispersal), and A = 10 X^2^/T (strong optimal dispersal). We ran each simulation for a period of time long enough that the initial transient states passed. We used the computed curves near the end of each simulation to measure approximate speed and amplitude (peak value) of the traveling waves of population density, as well as maximum trait variance attained at the population’s center. The results are shown in Figure 2. At extreme (unrealistic) gradients, optimal dispersal greatly enhances population density and lowers both (local) trait variance and extinction risk (Figure 2a). Similarly, a population’s expansion at extremely steep gradients is much faster when the individuals disperse optimally. The intraspecific trait variance is also controlled to much lower and more reasonable levels by the assortment effects of optimal dispersal at extreme gradients. Note that the relatively sharp decline in the amplitude of the population density waves at extreme gradients is predominantly due to the phenotypic load 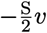 in (10c) which increases as *v* increases with gradient. That is why significantly slower decline in wave amplitude is observed when optimal dispersal is strong and effectively controls the rise in trait variance to more moderate values.

We note, however, that the large effects of optimal dispersal observed in Figure 2a occur mainly in extreme environmental gradients. Based on our previous discussions of plausible ranges of values of environmental gradients (Shirani and Miller, 2022: Sect. 3.2), such improvements mainly occur at exceedingly steep gradients which are unlikely to be biologically realistic. As the insets in the graphs of Figure 2a show, when the gradient in trait optimum takes more reasonable values between 0 and 2 Q/X, the improvements are much less pronounced. We observe significant changes in the expansion speed only at steep gradients greater than 1 Q/X. At more typically observed gradients below 1 Q/X, phenotype-optimal dispersal still facilitates local adaptation and significantly reduces within-population trait variance, but such effects appear to be less consequential for the population’s range expansion capacity.

In a typically shallow environmental gradient of ∇_*x*_Q = 0.2 Q/X with only diffusive dispersal (A = 0 X^2^/T), increased specialization raises population density and trait variance (Figure 2b). With optimal dispersal, both these quantities are regulated to lower levels. By contrast, Figure 2b shows that phenotype-optimal dispersal does not significantly increase range expansion speed, even when the phenotype utilization variance in the population is small. To be more precise, in the absence of phenotype-optimal dispersal the competitive release afforded by small values of V results in significant increase in both population density and trait variance as V → 0. The directed gene flow created by optimal dispersal, however, substantially depresses the effects of this competitive release. It controls the level of increase in trait variance to much lower values. As a result, competition remains strong even though small values of V tend to release the individuals from competition. When optimal dispersal propensity is very strong (A = 10 X^2^/T,) the effect of competitive release is fully depressed and the amplitude of population density waves remains close to the carrying capacity K = 1 N/X for all values of V. When *n* is controlled to almost constant values, decreasing V in the first (competition) term in (12f) to zero will eliminate the inflating effects of competition on trait variance. As a result, with strong optimal dispersal, steady-state trait variance will converge to a value mainly controlled by random gene flow (term (12c) in our model) and mutation-section balance (term −S*v*^2^ +U in (12f)). Since both U and the environmental gradient are relatively small in Figure 2b, this value is relatively small. This explains the sharp decline we observe in the curves of trait variance in Figure 2b as V → 0, both for A = 10 X^2^/T and for A = 4 X^2^/T.

Finally, we note that our numerically obtained traveling wave amplitudes and trait variances are close to values that can be obtained analytically. The curves of wave amplitude and trait variance in Figure 2 were computed approximately, by running the simulations for a sufficiently long time and measuring the (almost steady) values of these quantities at the center of the population at the end of the simulation. These curves can more accurately be computed by solving the equations of the fully-adapted spatially homogeneous (except possibly at the close vicinity of the habitat boundary) equilibrium of the model, similar to our computations in Section 4.2 of our previous work (Shirani and Miller, 2022). To perform such analysis—which importantly allows us to also derive equations for critical environment gradients—we first note that the range dynamics shown in Figure 1 suggests that, as *t* → ∞, the solutions eventually rich an equilibrium. Except in the close vicinity of the habitat boundary (where we modify the perceived gradient as described in Appendix A), the population density and trait variance at this equilibrium are spatially homogeneous and trait mean is equal to Q. We denote this equilibrium state by (*n*\*, *q*\*, *v*\*), where *n*\* *>* 0 and *v*\* ≥ 0. Sufficiently far from the habitat boundary, we have ∇_*x*_*n*\* = 0, *q*\* = Q, and ∇_*x*_*v*\* = 0. Since we assume Q to be linear, we further have 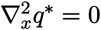.

From (10)–(12) we obtain, after some algebraic manipulations, that

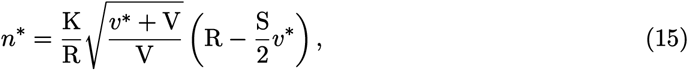

and

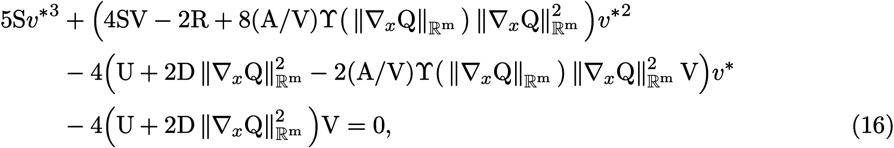

where 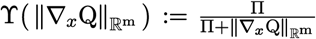, as in (9). The cubic algebraic equation (16) has a positive root for all nonzero values of 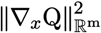. The graph of this root with respect to changes in the magnitude of the gradient 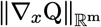 is approximately a straight line with positive slope, as shown in supplementary Figure S4a. Substituting this root into (15) for different values of 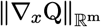 provides a graph of *n*\* with respect to 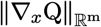. With the parameter values associated with Figure 2, this graph gives an approximate curve of wave amplitudes for each values of A, as shown in supplementary Figure S4a. We observe very close agreement between the approximate curves shown in Figure 2 and the accurate ones shown in Figure S4.

#### Remark 4.1 (Critical environmental gradients)

The results shown in Figure (2a) imply that, for every value of A, there exists a *critical environmental gradient* magnitude at which the curves of wave amplitude and wave speed concurrently vanish to zero. That is, the population fails to survive beyond the critical gradient—unless possibly marginally (with low density over a short range) at the vicinity of the habitat boundary. The equilibrium equations (15) and (16) allow us to derive a fairly simple equation for calculating the critical gradients. Setting *n*\* = 0 in (15) gives *v*\* = 2R/S, which is a solution of (16) when the environmental gradient satisfies

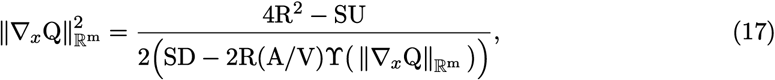

where 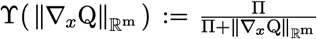 as we stated before. The solution to (17) gives the critical gradient magnitude, which we denote by 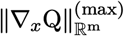. When dispersal is only random, that is A = 0, the critical gradient can be calculated as

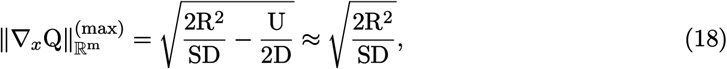

which is the same formula we derived in our previous work (Shirani and Miller, 2022: Remark 4). The approximation in (18) is made based on the fact that, typically, U ≪ 4R^2^/S.

#### Remark 4.2 (Slow growth rate, strong selection, and chance of survival)

The critical gradient magnitude 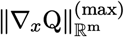 can be used to predict the chance of survival of a species under steep environmental stress gradients. Larger values of 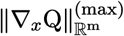 imply higher chance of survival (lower chance of extinction). In the absence of optimal dispersal, the critical gradient (18) implies that slowly-growing species (small R) and species under strong selection (large S) are at higher risk of extinction. To see if phenotype-optimal dispersal is sufficiently effective in increasing the survival chance of such species, we use (17) to compute critical gradients for species with relative slow growth rate (R *<* 1 T^*−*1^) and species under relatively strong selection (S *>* 0.5 Q^*−*2^/T). The results are shown in Figure 3. We first note that, in the absence of optimal dispersal A = 0 Q/X, these vulnerable classes of species are indeed at high risk of extinction within plausible ranges of values for 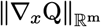 (that is, between 0 and 2 Q/X based on our estimates (Shirani and Miller, 2022: Sect. 3.2)). Phenotype-optimal dispersal appears to substantially increase the survival chance of the species. With strong optimal dispersal A = 10 Q*/tX*, for example, the species is unlikely to go extinct at plausible gradients (of course, excluding physical barriers), even with a very slow growth rate such as R = 0.2 T^*−*1^, or under an exceedingly strong selection such as S = 2 Q^*−*2^/T.

**Figure 3:**
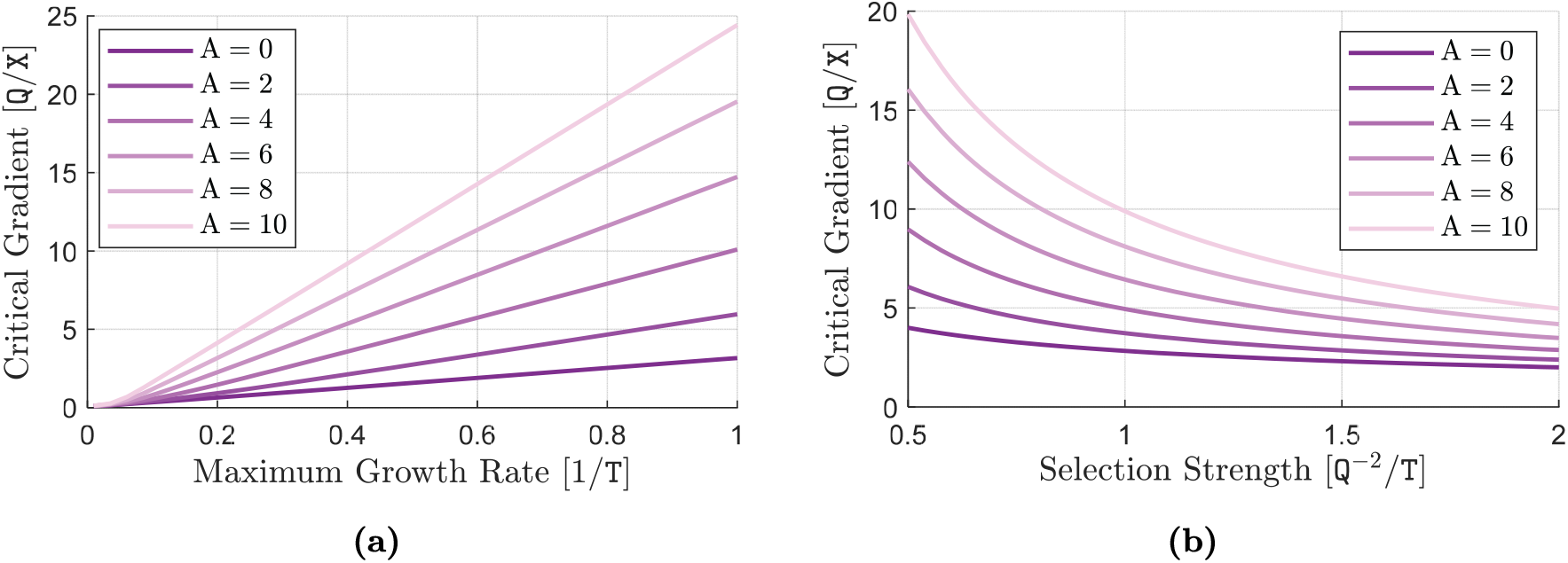
Effects of optimal dispersal on survival chance of slowly-growing species and species under strong selection. The critical environmental gradient 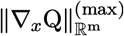 is computed by solving (17) and is plotted in (a) versus the maximum growth rate R, and in (b) versus the strength of stabilizing selection S. The larger the critical gradient the higher the chance of survival of the species in steep environmental stress gradients. In each graph, computations are performed for six different values of A in the range from 0 X^2^/T to 10 X^2^/T. Parameter R takes relatively small values in (a) and the constant value R = 2 T^*−*1^ in (b). Parameter S takes the constant value S = 0.2 Q^*−*2^/T in (a) and relatively large values in (b). The rest of the model parameters take their typical values given in Table 1.

### 4.3 Phenotype-Optimal Dispersal Enhances Local Adaptation at Range Margins

We now study more specifically how the mean adaptation rate, *∂*_*t*_*q*, is affected by random and non-random (directed) components of gene flow, particularly at range margins (wavefronts). The sum of the terms (11a) and (11b) represents the contribution of random gene flow in determining the rate of change of trait mean, whereas the sum of the terms (11c)–(11e) presents the contribution of directed gene flow. The sum of all terms (11a)–(11e) determines how the trait mean is changed due to individuals’ dispersal, both random and directed. We illustrate the effects of each of these three contributions separately, using the same simulation layout as used for the results shown in Figure 1b, and the same solutions of the model that we computed there. The resulting curves as the population expands its range over time are shown in Figure 4. Due to symmetry, the curves are shown only for the right half of the habitat. Positive values in each curve at a point *x* imply that the corresponding component of gene flow represented by the curve tends to increase the trait mean *q* at *x*. Contrariwise, negative values imply a tendency to decrease *q*. Since the initial profile of trait mean at *t* = 0 T, as shown in Figure 1b, is below the trait optimum Q over the right half of the habitat, positive values of the curves imply adaptive effects (increasing *q* toward Q), and negative values imply maladaptive effects.

**Figure 4:**
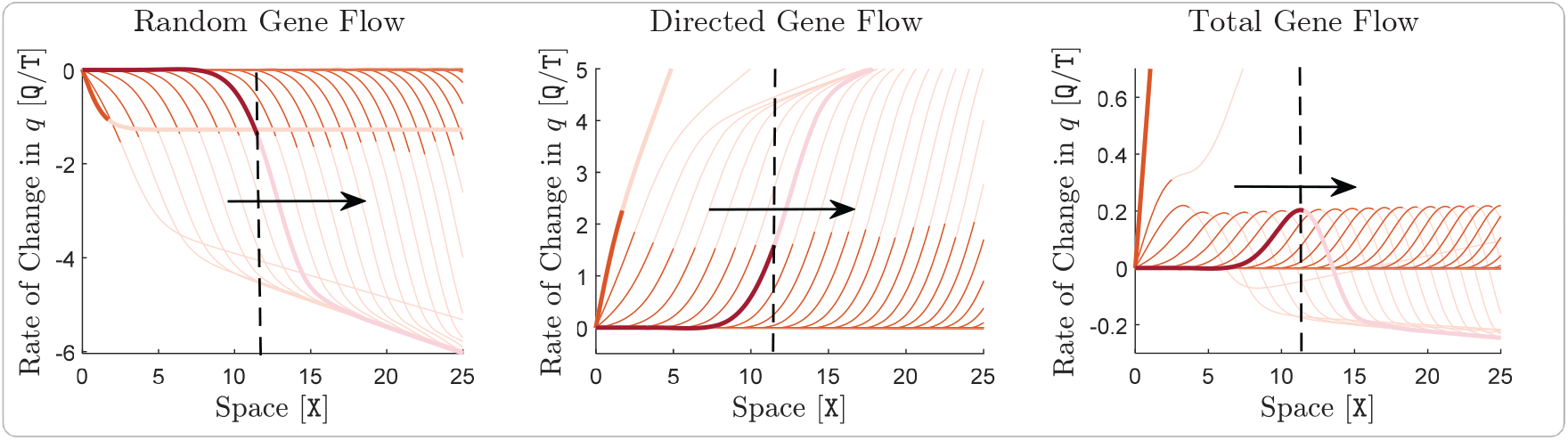
Effects of gene flow on local adaption of a species. Contribution of random gene flow to rate of change of the trait mean, *∂*_*t*_*q*, is shown on the left. The curves of this graph are computed as the sum of the terms (11a) and (11b), which capture the effects of random dispersal on *∂*_*t*_*q*. In the middle, contribution of directed gene flow to *∂*_*t*_*q* is shown. The directed gene flow is generated by optimal dispersal and its contribution is computed as the sum of the terms (11c)– (11e). Contribution of the total (net) gene flow to *∂*_*t*_*q*, that is, the sum of the curves shown in the middle and the left graphs, is shown on the right. The graphs correspond to the same simulation of a species’ range dynamics as in Figure 1b, that is, when ∇_*x*_Q = 1.5 Q/X and A = 10 X^2^/T. In all graphs, the computed curves are shown only on the right half of the habitat. The curves extend symmetrically about the origin to the left half of the habitat. Moreover, the portion of each curve that lies outside the effective range of the species, that means over the regions where the population density rapidly vanishes to zero, has been made transparent. In all graphs, curves are shown at every 1 T. The descriptions of the highlighted curves, arrows, and dashed lines are the same as those provided in Figure 1b.

Gene flow to range margins (wavefronts) induced by phenotype-optimal dispersal is always adaptive, but is of little consequence except at steep environmental gradients. At such gradients, optimal dispersal speeds up adaptation at both the range center and range margins, although a perfect match of trait mean and trait optimum is never achieved at the wavefront. To see this, we first note that Figure 1b showed rapid convergence of *q* to Q at the population’s core, due to the strong effect of phenotype-optimal dispersal at the steep gradient ∇_*x*_Q = 1.5 Q/X. This rapid convergence can also be clearly observed in Figure 4. The rate of change in *q* quickly approaches zero at the core of the population after a couple of generations. At range margins, however, Figure 1b shows that adaptation never occurs perfectly. The core-to-edge random gene flow created by random dispersal is always maladaptive, whereas the directed gene flow generated by phenotype-optimal dispersal is always adaptive. Importantly, the total gene flow to range margins appears to always be adaptive, implying that phenotype-optimal dispersal not only compensates for the maladaptive effects of random movements, but also reverses their effects on local adaptation of the marginal populations. Similar observations apply when we analyze the adaptive/maladaptive effects of different components of gene flow associated with the simulation results shown in supplementary Figure S3b, that is, when the environmental gradient is shallow (∇_*x*_Q = 0.2 Q/X) but individuals are highly specialized (V = 1 Q^2^). The resulting curves are shown in Figure S5. The difference in this case—possibly because of the complicated interaction between the effects of optimal dispersal and competition—is that convergence at the core shows overshooting dynamics, during which the curves of directed gene flow take negative values (but still adaptive) to decrease the overshot *q* back to Q.

Total gene flow to the wavefront remained adaptive in further simulations with shallower (i.e. more typical) environmental gradients and lower degrees of specialization (larger V). In these simulations, we repeated our analysis like that shown in Figures 4 and S5 for different slopes of the environmental gradient and different values of V. The sample curves in Figure 4 show that the maximum adaptive (or maladaptive) effects of directed (or random) gene flow occur at the edge of the population. Therefore, we used the value of the total contribution of gene flow to *∂*_*t*_*q*, computed at the edge of the population, as our reference for measuring the significance of phenotype-optimal dispersal in facilitating adaptation at range margins. The results are shown in Figure 5. We see that the total gene flow remains adaptive to the range margins, even when the environmental gradient is fairly shallow or individuals are generalists. This implies that phenotype-optimal dispersal is quite effective in compensating for the maladaptive effects of core-to-edge random gene flow. Yet the curves shown in Figure 5, along with our previous observations through Figure 2, suggest that the adaption facilitated by phenotype-optimal dispersal at range margins sufficiently enhances range expansion capacity of the population primarily when environmental gradients are steep (slopes greater than 1 Q/X).

**Figure 5:**
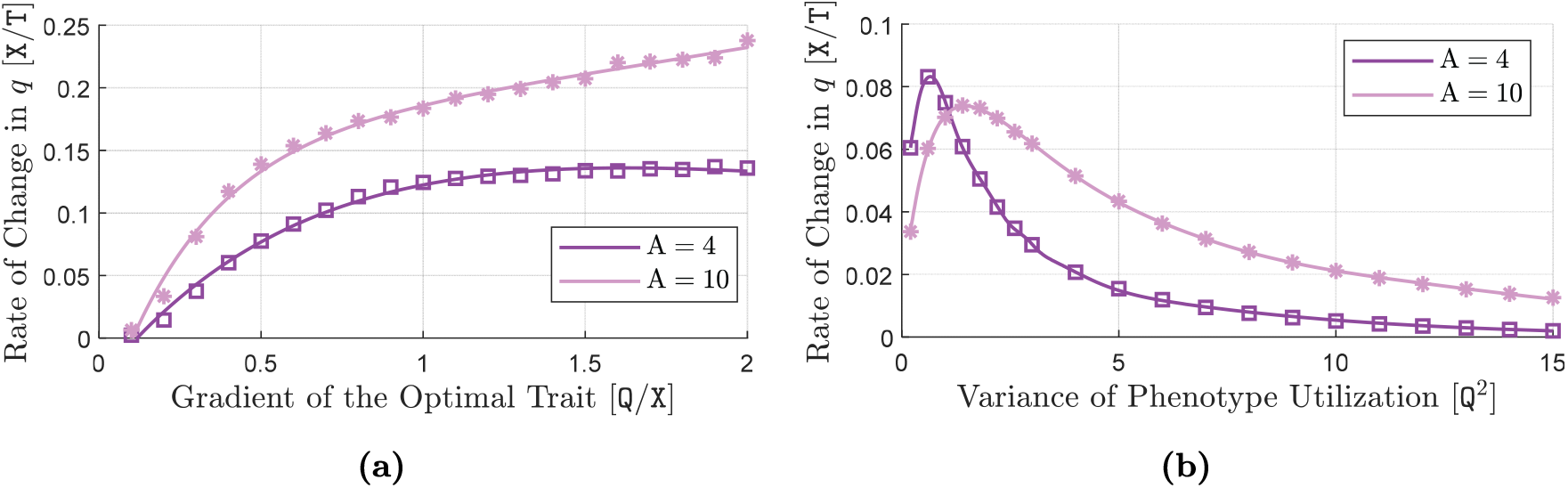
Effects of directed gene flow on local adaption at the edge of a species’ range. The contribution of total (net) gene flow to rate of change of the trait mean, *∂*_*t*_*q*, at the edge of a species’ range is shown in each graph, for two different values of A. In (a), the species’ phenotype utilization variance takes its typical value of V = 4 Q^2^, and the magnitude of ∇_*x*_Q is made variable. In (b), the environmental gradient takes its typical value of ∇_*x*_Q = 0.2 Q/X, and V is made variable. The rest of the model parameters take their typical values given in Table 1, with m = 1. The data points in graphs (a) and (b) are obtained as follows. At each value of ∇_*x*_Q in (a) and each value of V in (b), a simulation is performed for a period of time long enough that the species’ range expansion dynamics reaches an approximate steady-state. In each simulation, the contribution of the total gene flow to *∂*_*t*_*q* is computed for the right half of the species’ range, as the sum of all terms (11a)–(11e) in (11). The graphs shown on the right of Figures 4 and S5 show samples of such computation results. The value of the total gene flow contribution that is obtained at the edge of the species’ range at the end of each simulation is then shown in graphs (a) and (b).

We note that the curves associated with A = 0 are not shown in Figure 5. In fact, with A = 0 we only have random gene flow (random dispersal), the effect of which is already known to be maladaptive at any nonzero values of the environmental gradient. This means that the curves for A = 0 in Figure 5 would take negative values for all environmental gradients and utilization variances. The larger the gradient the more maladaptive effect of random gene flow.

### 4.4 Phenotype-Optimal Dispersal Promotes Persistence Under Abrupt Environmental Fluctuations

We carried out simulations with periodic abrupt environmental shifts, to test the prediction that phenotype-optimal dispersal could improve population survival under such shifts. Rapid adaptation of individuals within a single generation, as facilitated by adaptive dispersal strategies such as matching habitat choice, is predicted to be crucial to the survival of a population under climate change, particularly when changes are sharp and frequent (Bonte et al., 2012; Edelaar and Bolnick, 2019; Jacob et al., 2017; Nicolaus and Edelaar, 2018). To explore how phenotype-optimal dispersal will affect the range dynamics of a species under abrupt climatic changes, we simulate our model in a one-dimensional habitat with a steep trait optimum gradient of ∇_*x*_Q = 1.5 Q/X, wherein in the trait optimum periodically fluctuates (shifts) up and down with no change in its gradient. For this, we initialize our simulation at *t* = 0 T with a relatively established population at the center of the habitat. This initial population is obtained as the curves at *t* = 4 T of a preliminary simulation similar to the one shown in Figure 1, with 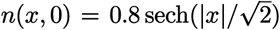, *q*(0, 0) = Q(0), ∇_*x*_*q*(*x*, 0) = 0.7 ∇_*x*_Q, and *v*(*x*, 0) = 1 Q^2^. The thick orange curves shown in Figure 6 indicate the initial populations. To simulate abrupt temporal fluctuations in the environment, we uniformly shift up the line of trait optimum Q by a certain fluctuation amplitude at the beginning of a fluctuation period, and then shift it back down by the same amplitude at the middle of the period. We repeat these fluctuations periodically, starting at *t* = 0 T, with a relatively short period of 2 T. With purely random dispersal plus environmental shifts, population density fluctuates, but is always significantly lower than in a constant environment. Figure 6a shows the simulation results for an environmental fluctuation amplitude of 5 Q, when dispersal is only random, A = 0 X^2^/T. The high level of maladaptation that is abruptly induced in the population at *t* = 0 T, when the optimum gradient is shifted up, quickly reduces both population density and and expansion speed. Yet directed natural selection acts on the large deviation of trait mean from the trait optimum, and the population gradually adapts to the new environmental optimum. However, since the period of the fluctuations is relatively short, the population will not fully recover its peak density before experiencing another abrupt change in the environment. The periodically repeated loss-and-recovery dynamics of the population density eventually reach a steady-state, at which the peak population density fluctuates between a fixed high and a fixed low value as the populations expands its range. The red and blue curves in Figure 6a show samples of population density profiles at such high and low extremes.

**Figure 6:**
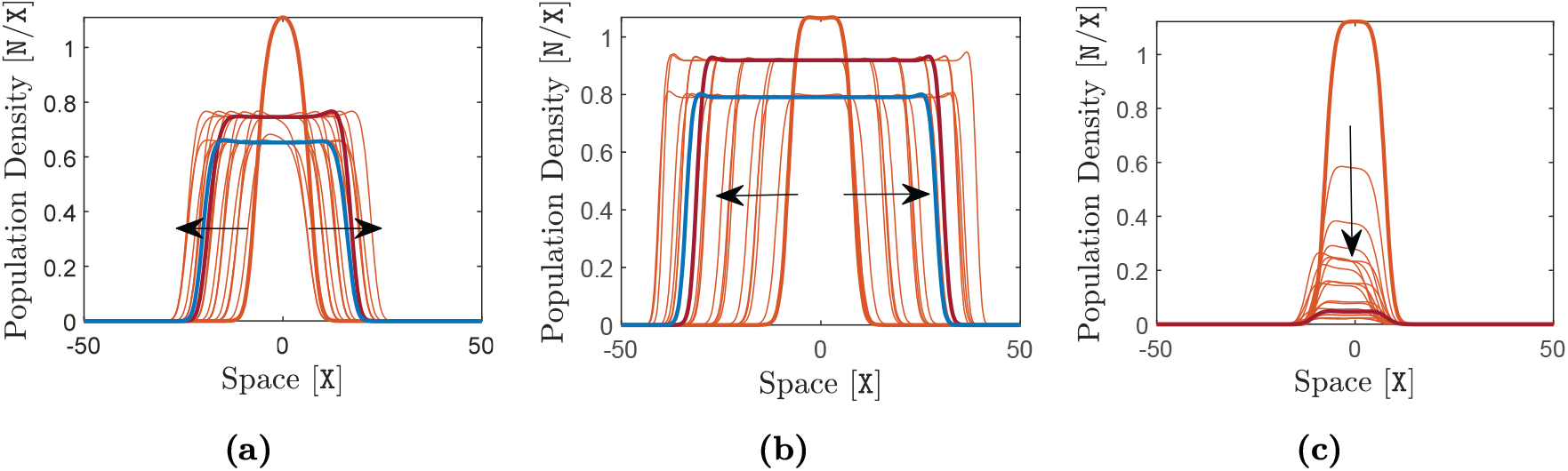
Range dynamics of a species under periodic abrupt fluctuations in the environmental trait optimum. Here, *m* = 1, ∇_*x*_Q = 1.5 Q/X, and A takes different values in each graph. The rest of the model parameters take their typical values given in Table 1. Graph shows range dynamics without optimal dispersal, that is A = 0 X^2^/T. Graph (b) shows range dynamics with strong optimal dispersal, A = 10 X^2^/T. Graph (c) shows extinction of a species with the typical value A = 4 X^2^/T due to high-amplitude environmental fluctuations. In all graphs, the period of abrupt fluctuations in the trait optimum is 2 T. At the beginning of each period, the trait optimum Q is shifted up by a preset fluctuation amplitude and remains at this value for the first half of the period. Then, it is shifted down by the same amplitude to the initial value and remains at this value for the second half of the period. The fluctuation amplitude is set equal to 5 Q in (a) and (b), and equal to 9 Q in (c). The thick orange curves indicate the initial curves at *t* = 0 T and arrows show the direction of evolution in time. In (a) and (b), curves are shown at every 1.5 T, and two sample curves are highlighted at *t* = 20 T (in red) and *t* = 20.5 T (in blue). In (c), curves are shown at 20 logarithmically distributed time samples, with the first curve after the initial curve being shown at *t* = 0.1 T. A sample curve at *t* = 20 T is also highlighted in red.

Strong phenotype-optimal dispersal, A = 10 X^2^/T, enhances both range expansion speed and steady-state population density under the abrupt environmental fluctuations that we simulate here. The results are shown in Figure 6b. This is because the large phenotype-environment mismatch perceived by individuals, immediately after an abrupt shift in the optimum phenotype, creates a strong phenotypic potential for the individuals to disperse to better-matching locations. As a result, the population can rapidly adapt to the new environmental optimum when a change occurs, and hence it loses much less of its density. However, in agreement with our observations in the previous studies described above, the significant effects of optimal dispersal that we observe here are in the presence of a steep environmental gradient of ∇_*x*_Q = 1.5 Q/X. In shallower (more typical) gradients, as the supplementary Figure S6 shows, the effects are much less noteworthy.

When environmental fluctuations are very large in amplitude, phenotype-optimal dispersal can allow survival when a purely diffusing population goes extinct. When the environmental shifts are too large, the density loss due to the excessive level of maladaptation after each shift will be too high to be fully recovered by the adaptation that occurs afterwards. As a result, the population will not be able to reach a persistent state and its density keeps decreasing due to the repeated changes in the environment. Eventually, the population becomes extinct. Figure 6c shows an example of such extinction dynamics, when phenotype-optimal dispersal is still fairly strong, A = 4 X^2^/T, but environmental fluctuations occur with a large amplitude of 9 Q. To see if phenotype-optimal dispersal increases the chance of survival when amplitude of the fluctuations is large, we repeat the simulation associated with Figure 6 for different values of fluctuation amplitudes. For each value, we compute the average value of the fluctuations in population density when it reaches a steady-state, and show it in Figure 7a. We see that when the amplitude of the environmental fluctuations is not very large, optimal dispersal does not significantly increase the sustained density of the population. At very large fluctuation amplitudes, however, optimal dispersal can greatly increase the survival chance of the species.

**Figure 7:**
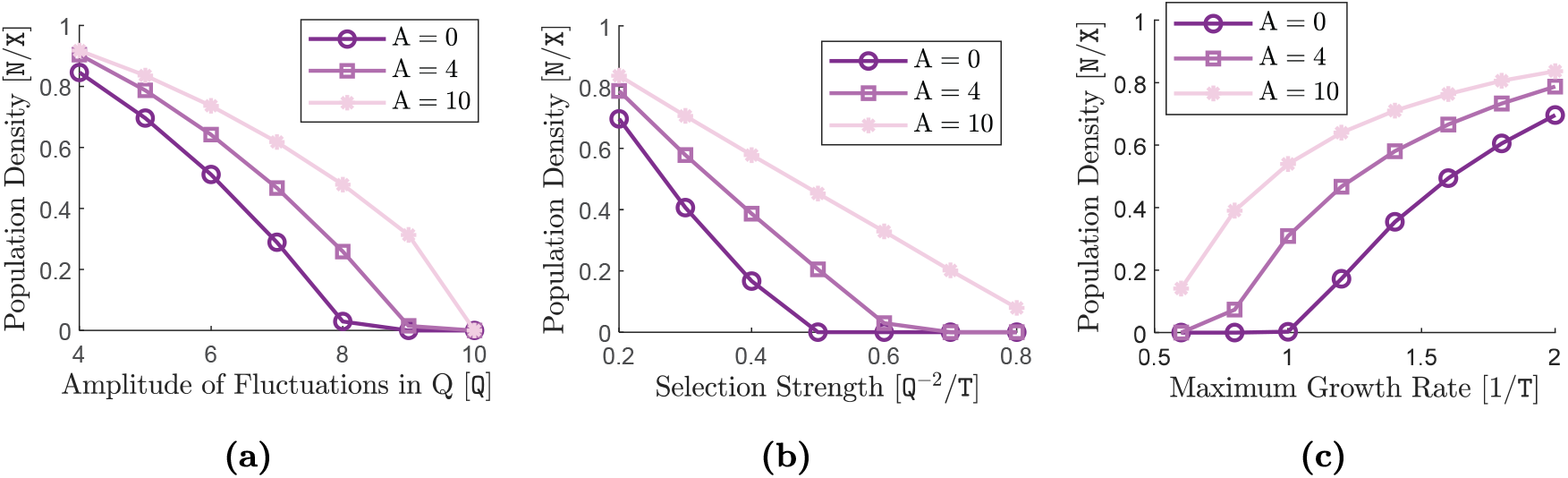
Steady-state mean population density of a species under periodic abrupt fluctuations in the environmental trait optimum. Here, *m* = 1, ∇_*x*_Q = 1.5 Q/X, and A takes different values in each graph. The same periodic abrupt fluctuations as described in Figure 6 is applied to the trait optimum Q, with the typical amplitude of 5 Q in (b) and (c) and variable amplitude in (a). Except for S and R which are made variable in (b) and (c), respectively, the rest of the model parameters take their typical values given in Table 1. At each value of the variable parameter shown at the horizontal axis of each graph, and each value od A, the simulation is run for a sufficiently long period of time so that the amplitude of the fluctuations in population density of the species reaches a steady-state. The minimum and maximum values of such periodic fluctuations (peaks of the blue and red curves as shown in Figures 6a and 6b) are calculated near the end of the simulation, and their average value is shown by different markers for each value of A, along with an interpolated solid line. In (a), the steady-state mean value of the population density is shown with respect to changes in the amplitude of the abrupt fluctuations in Q. In (b), the steady-state mean value of the population density is shown with respect to changes in the strength of stabilization selection, S. In (c), the steady-state mean value of the population density is shown with respect to changes in the maximum growth rate of the species, R.

Varying the strength S of natural selection shows that optimal dispersal increases mean population density markedly only under very strong selection. Strong natural selection amplifies the effects of maladaptation induced by abrupt environmental changes, as implied from the term 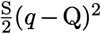 in (10c). As a result, when S is large, the population suffers from a greater density loss after each shift in the trait optimum. However, larger values of S expedite adaptation to the new environment, due to (11f). To see how these conflicting effects of stronger natural selection affect population density under environmental fluctuations, and whether or not phenotype-optimal dispersal can be sufficiently advantageous to ensure population survival, we repeat the simulation associated with Figure 6 for different values of S and measure the steady-state average value of the fluctuations in population density. The results are shown in Figure 7b. In general, we see that stronger natural selection reduces the sustained population density under the abrupt fluctuations we simulated. When stabilizing section is weak, as we typically observe in nature (Kingsolver et al., 2001), phenotype-optimal dispersal does not substantially increase mean population density. However, under very strong stabilizing selection, we observe that the population’s chance of survival is substantially increased if its individuals disperse optimally.

Phenotype-optimal dispersal can allow a population with a low maximum intrinsic growth rate R to persist under environmental fluctuations that would drive it to extinction if it carried out only diffusive dispersal. The maximum growth rate of the population is a key factor in accelerating population density recovery following density losses caused by environmental changes. Slowly growing populations will have a lower chance of recovering their full density before suffering another loss. To see how this impacts the population survival, we repeat our simulations for different values of R and show the steady-state average value of population density fluctuations in Figure 7c. We see that slowly-growing populations maintain a significantly lower density under environmental fluctuations, compared with fast-growing populations. However, phenotype-optimal dispersal substantially increases the survival prospects of slowly-growing species. In particular, with the relatively large fluctuation amplitude of 5 Q that we considered in our simulation, we observe that, when R takes a relatively low value of approximately 1 T^*−*1^, a randomly dispersing population becomes extinct whereas a population with strong optimal dispersal can persist. We note that, with our choice of generation time as the unit of time T, the maximum intrinsic growth rate is argued to be a demographic invariant within some homogeneous taxonomic groups (Niel and Lebreton, 2005). In particular, a slow growth rate of R = 1 T^*−*1^ is observed in a variety of taxa such as birds, sharks, and mammals (Dillingham et al., 2016; Niel and Lebreton, 2005; Shirani and Miller, 2022), which are often sufficiently mobile to evolve optimal dispersal strategies.

### 4.5 Phenotype-Optimal Dispersal in Fragmented Habitats

We next ask whether optimal dispersal can enhance survival and range expansion speed when the habitat is fragmented. In highly fragmented environments, the cost of randomly moving from one habitat patch to reach another patch is particularly high. Optimal habitat selection can also be harder in such environments, as optimal detection of matching habitat patches can be substantially imperfect and dispersal mortality can be higher (Cote et al., 2017). However, the enhanced local adaptation facilitated by optimal dispersal can still be imagined to be sufficiently beneficial to compensate for such costs. Therefore, it is argued that habitat choice behavior and adaptive dispersal strategies should also be selected in fragmented landscapes, and might be essential for the persistence of populations in such environments (Bonte et al., 2012; Cote et al., 2017). Here, we investigate whether or not phenotype-optimal dispersal can be of particular importance to a population’s adaptation and range expansion capacity when the available habitat is highly fragmented. We first specify our simulated fragmented environment. We consider a two-dimensional habitat

Ω = (−50, 50) × (−50, 50), with a trait optimum profile that changes linearly in the *x*_1_−direction (horizontal axis) and is constant in the *x*_2_−direction (vertical axis). We set a relatively steep gradient of 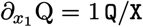 in *x*_1_−direction. Since habitat loss directly affects the carrying capacity of the environment (Baguette et al., 2013), we simulate habitat fragmentation by setting a patchy profile for the carrying capacity parameter K(*x*). This profile is shown in supplementary Figure S7, and its specific pattern can also be approximately seen through the last frames (at *t* = 40 T) in Figure 8. If patch sizes are large in both directions, relative to the average (random) dispersal distance per generation, the range dynamics inside each patch is expected to be similar to that in the continuous habitat cases we studied before. Therefore, to make the effect of fragmentation sufficiently pronounced, we consider relatively narrow patches of width 2 X—noting that the random dispersal coefficient D takes the typical value of 1*I*_2_ X^2^/T, where *I*_2_ denotes the 2×2 identity matrix.

**Figure 8:**
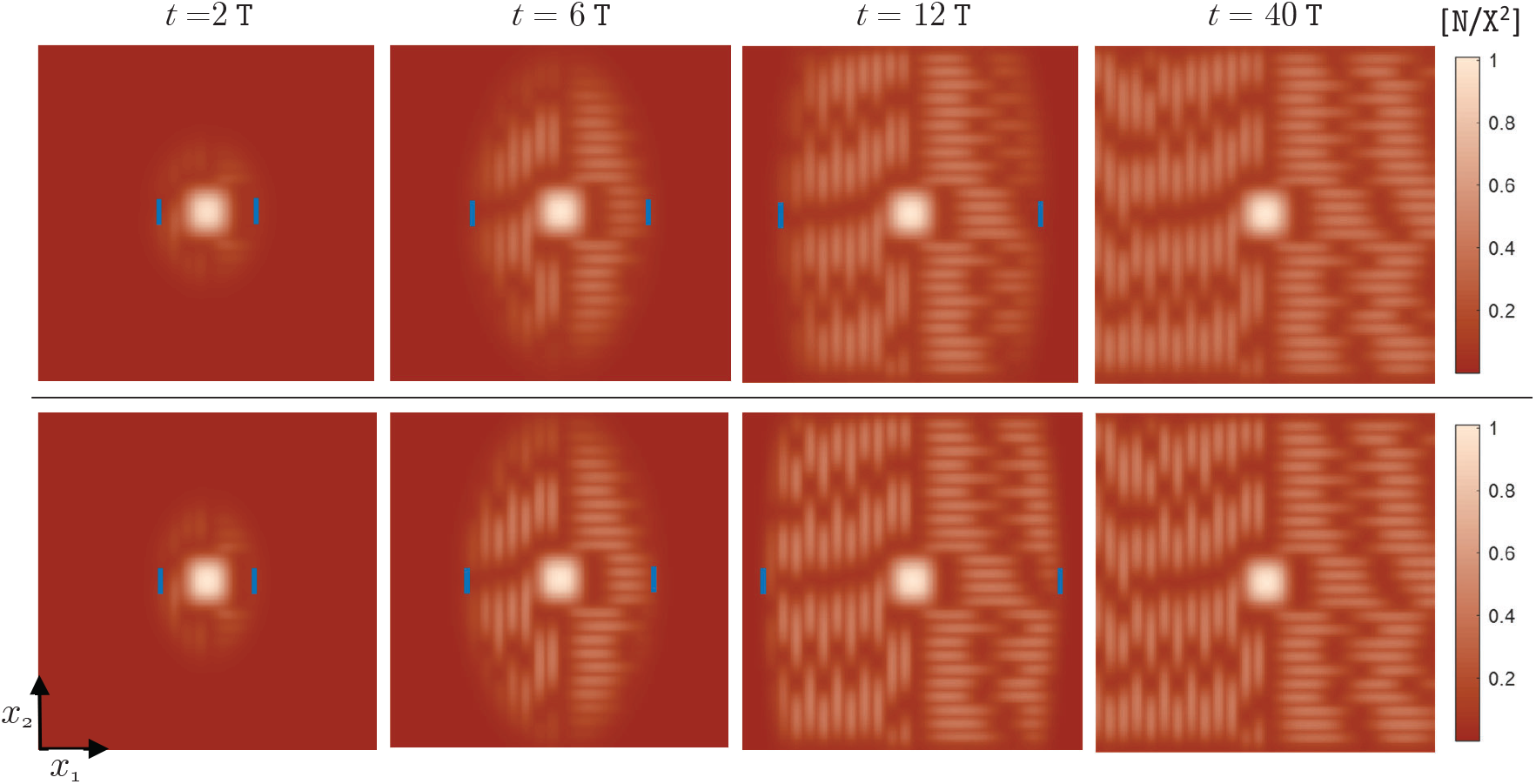
Range expansion of a species in a fragmented two-dimensional habitat. Here, m = 2, the environmental gradient along the *x*_2_-axis is zero, and the environmental gradient along the *x*_1_-axis is 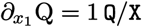. Habitat fragmentation is simulated by considering a patchy profile for the carrying capacity K, as shown in the supplementary Figure S7. Parameter A takes different values for the results shown in the upper and lower panels. The rest of the parameters take their typical values given in Table 1. Four frames of the spatial profile of the species’ population density are shown in each panel as the species’ range evolves in time. The upper panel shows range expansion of a species with no optimal dispersal, A = 0 X^2^/T, whereas the lower panel shows the range expansion of the species with strong optimal dispersal, A = 10 X^2^/T. For comparison purposes, the same simulations but with constant carrying capacity of K = 1 N/X^2^ are performed and the leftmost and rightmost edges of the population are indicated by blue bars in each plot. It should also be noted that the final (approximate steady-steady) profile of the phenotype-environment mismatch and trait variance at *t* = 40 T are also shown in supplementary Figures S8 and S9.

We let the length of these rectangular patches take values between 10 X and 15 X. On the right half of the habitat, we arrange the patches horizontally, so that they are stretched in the direction of the environmental gradient. On the left half of the habitat, we arrange the patches vertically so that they are stretched perpendicular to the environmental gradient. We consider this specific layout to further observe if the alignments of the patches with the environmental gradient can have a particular impact on the range expansion dynamics. We arrange the rectangular patches side-by-side, with no gap between them. However, over each patch, we smoothly decrease the value of K from 1 N/X^2^ at the patch’s center to 0.05 N/X^2^ at the patch’s edge. This leaves fairly inhabitable regions of low carrying capacity between patch cores, as shown in Figure S7. Finally, we make an exception for the size of the patch located at the center of the habitat, and let it be large. We initialize the population in this central patch, so that it can get well-established before expanding through the highly fragmented areas.

When dispersal is entirely random, population density drops off significantly over the patches and mismatch between the trait mean and trait optimum grows sharply from the center to the edge of the patches. Figure 8 shows the simulation results for a species with only randomly dispersing individuals (upper panel) and for a species with phenotype-optimal dispersal (lower panel). To make a comparison with range expansion in a continuous habitat, we also perform the simulations with a constant carrying capacity of K = 1 N/X^2^ and indicate the leftmost and rightmost edges of the population by blue bars in Figure 8. The population establishes itself in the central patch to almost its maximum capacity of approximately 1 N/X^2^. However, it spreads over the fragmented regions with much lower density. This is mainly due to a significant level of maladaptation that is maintained by the homogenizing effect of random gene flow. Figure S8 shows the phenotype-environment mismatch (*q* −Q) at the end of the simulations, when the mismatch has approximately reached a steady state over the whole habitat. The steady-state level of mismatch is low at the center of the patches. However, it increases rather sharply towards their edges. Since patches are relatively narrow, this results in a relatively high level of overall maladaptation over a large area of the patches, and hence the significant loss of population density.

Phenotype-optimal dispersal raises the rate, though not the steady-state level, of adaptation within habitat patches. It thereby enhances the transient expansion dynamics of the population, as observed through the increased expansion speed in the lower panel of Figure 8. However, we observe that optimal dispersal does not significantly improve the steady-state level of adaptation (shown in Figure S8) and population density within the patches. The maximum population density that we observe in patches of the fragmented region at *t* = 40 T is approximately 0.45 N/X^2^ when A = 10 X^2^/T, which is only slightly larger than the maximum density of 0.41 N/X^2^ observed when A = 0 X^2^/T. This is because, due to the narrow widths of the patches, the directed gene flow created by phenotype-optimal dispersal is not sufficiently strong to effectively compensate for the maladaptive core-to-edge random gene flow within the patches.

With phenotype-optimal dispersal, the rate of range expansion in the simulated fragmented habitat almost equals that in a continuous habitat; although the population density in the fragmented habitat remains significantly lower. This is implied by the locations of the blue bars in Figure 8, which also coincide with the leftmost and rightmost edges of the population in the fragmented habitat. Notably, we do not observe any effects of phenotype-optimal dispersal that particularly enhance the range expansion capacity of the population when a habitat is fragmented, compared with the continuous habitat case.

With phenotype-optimal dispersal, the maximum steady-state (*t* = 40 T) population density on vertical patches is about 10 percent higher than the maximum density on horizontal patches. By contrast, when dispersal is only random, the particular layout of the patches—horizontally arranged on the right and vertically arranged on the left—does not result in any noticeable asymmetry in population’s expansion to each side of the habitat. Slightly better adaptation and lower trait variance are also observed under optimal dispersal within vertical patches. This can be explained by noting that the vertical patches in our simulation are stretched perpendicular to the gradient of the trait optimum. Therefore, optimal movements in the direction of the environmental gradient lead to a distribution of relatively close phenotypes along each of the vertical patches. Since the patches are also relatively narrow, the overall closeness of phenotypes in vertical patches decreases the maladaptive impacts of the random component of gene flow within the patches. As a result, population density can grow to (somewhat) higher levels within vertical patches.

Finally, in fragmented as well as in continuous habitat, phenotype-optimal dispersal markedly reduces trait variation within the patches. Figure S9 shows the final profiles of trait variance at *t* = 40 T. In the presence of a steep environmental gradient, as we consider here, the reduced phenotypic variability within patches results in substantial increase in variability among the patches. Although not included in our model, it can be argued that this enhanced genetic differentiation between the patches contributes to reproductive isolation between the local populations. Large genetic differences between the patches can reduce the propensity of individuals to move between the patches, to avoid mating with individuals in a genetically novel population (Benkman, 2017; Bolnick and Otto, 2013; Cote et al., 2017; Garant et al., 2007). The reduced movements between the patches and reduced genetic variation within the patches can then promote the evolution of assortative mating, which further reinforces the reproductive isolation between the patches. This process can eventually lead to sympatric speciation in the habitat (Bolnick and Otto, 2013; Garant et al., 2007; Lenormand, 2002; Nicolaus and Edelaar, 2018).

## 5. Discussion

Matching habitat choice has been hypothesized to have distinctive evolutionary effects on species’ range dynamics and genetic structure (Bolnick and Otto, 2013; Edelaar and Bolnick, 2019; Edelaar et al., 2008; Jacob et al., 2017; Nicolaus and Edelaar, 2018). Because of the potentially complex interactions between matching habitat choice and other eco-evolutionary forces, intuitive or verbal predictions of its effects must be taken skeptically. We therefore proposed and numerically studied a deterministic PDE model incorporating a specific type of matching habitat choice we called phenotype-optimal dispersal, together with other eco-evolutionary forces. Our goal was to quantify the environmental and other conditions under which phenotype-optimal dispersal has substantial effects on the spatial dynamics of population density as well as of the mean and variance of a quantitative trait. To accomplish this, we compared the results of simulations incorporating a combination of phenotype-optimal and random (diffusive) dispersal with the results of simulations incorporating random dispersal only.

A key novel aspect of our model, among studies of matching habitat choice, is that it is a deterministic continuum model, particularly suitable for modeling populations that disperse over (mostly) nonfragmented habitat. We also note that the strength of phenotype-optimal dispersal in our model depends on both phenotype-environment mismatch and environmental gradient. Dependence only on the phenotype-environment mismatch can result in redundant movements in shallow gradients which may be unlikely to evolve as a matching habitat choice strategy in nature. Our model also incorporates phenotype-dependent competition. Competition tends to inflate phenotypic variance. At the same time, matching habitat choice reduces local phenotypic variance, which should intensify competition. We used our model to study the nonintuitive range dynamics that results from the interaction between these two counteracting forces.

Through our extensive computational investigations, we found effects of phenotype-optimal dispersal on diverse measures of persistence, range expansion, adaptation and trait variance. These effects were generally important only in steep environmental gradients or when specialization was very pronounced, and in particular for slowly-growing species.

### 5.1 Implications for Adaptation, Range Dynamics, and Speciation

In a static environment with steep gradient, we found as expected that phenotype-optimal dispersal facilitates rapid adaptation of the individuals (and the population) within a single generation. To the extent that the relevant trait is heritable and the environmental gradient monotone, this should improve the fitness of subsequent generations and cause adaptive evolution. Optimal dispersal also lowers phenotypic variance both at central and peripheral populations, reducing further the need for natural selection to eliminate unfit phenotypes. In light of these adaptive effects, it is noteworthy that we found the gene flow created in the presence of optimal dispersal always to be adaptive; see Figures 4 and 5. Specifically, the directed gene flow induced by optimal dispersal effectively compensates for the maladaptive (swamping) effects of asymmetric random gene flow at range margins. As a result, we found that (in steep environmental gradients) optimal dispersal increased the speeds of traveling waves modeling range expansion; see Figures 1, and 2. It had little effect on peak population density behind the wavefront, but it significantly increased the growth rate of the population during transient states of population establishment. These effects also broadly persisted in fragmented habitats; see Figure 8.

Genetic swamping of peripheral populations by the gene flow from central populations has been hypothesized, for a long time, as a major cause of range limits—by destabilizing the migration-selection equilibrium at range margins (Haldane, 1956; Kirkpatrick and Barton, 1997; Lenormand, 2002; Mayr, 1963; Sexton et al., 2009). This hypothesis has been challenged by several theoretical and empirical studies (Case and Taper, 2000; Kottler et al., 2021; Shirani and Miller, 2022). We note that the present study also found no conditions—even in fragmented habitat—that led to a stable localized population (“range pinning”) in a linearly changing environment. Rather, populations either advanced without limit in a traveling wave or went extinct under violent environmental fluctuations. Indeed, our finding that gene flow to the wavefront was always adaptive provides further theoretical evidence that range pinning via genetic swamping should be rare in nature. This finding runs contrary to the prediction of Kirkpatrick and Barton (1997) based on a PDE model with constant variance, but consistent with the predictions of Barton (2001) with evolving variance, Case and Taper (2000) with competition and constant variance, and of the authors with competition and evolving variance (Shirani and Miller, 2022). It is also consistent with the earlier observation of Holt (1987) that habitat selection should often vitiate the swamping effects of gene flow as a constraint on the evolution of local adaptation. Empirical results reviewed by Kottler et al. (2021) have also found limited support for asymmetry in (total) center-to-edge gene flow, and very little evidence that such gene flow reduces mean fitness in edge populations.

We also note that rapid adaptation due to phenotype-optimal dispersal is particularly important in temporally changing environments, as it remarkably enhances population persistence under sharp environmental fluctuations; see Figures 6 and S6. This can be specifically important for slowly-growing species, such as birds and mammals, since they cannot recover sufficiently fast from population losses due to sharp changes in the environment; see Figure 7. Therefore, our results support existing arguments for taking matching habitat choice into account when predicting the response of species to climate-induced environmental fluctuations (Bonte et al., 2012; Edelaar and Bolnick, 2012, 2019; Jacob et al., 2017; Nicolaus and Edelaar, 2018; Pellerin et al., 2019).

It is worth noting that Pellerin et al. (2019) predict, based on their model, that climate change can result in contraction and fragmentation of the species’ range. By contrast, we do not see range contraction in our results even in the absence of phenotype-optimal dispersal (with pure random dispersal). Extinction by strong environmental fluctuations in our model is in the form of vanishing population density rather than contracting range. It is possible that the range contraction or fragmentation observed by Pellerin et al. (2019) could, to some extent, be due to the discrete patchy environment in their model and discrete movements of their individuals (they jump from one patch to the other at the end of each year). This provides an example, specifically in the context of matching habitat choice, of how range dynamics in continuous habitats can exhibit behavior unlike that in discrete habitat models.

Finally, we note that in fragmented habitats, the reduced within-population trait variation and enhanced local adaptation resulting from optimal dispersal increased the level of trait variation between local populations. Thus our results provide further quantitative theoretical support for the predictions that matching habitat choice can drive reproductive isolation, promote assortative mating, and contribute to sympatric or parapatric speciation (Berner and Thibert-Plante, 2015; Bolnick and Otto, 2013; Coyne, 1992; Edelaar et al., 2008, 2023; Endler, 1977; Holt, 1987; Kirkpatrick and Ravigné, 2002; Maynard Smith, 1966; Nicolaus and Edelaar, 2018; Ravigné et al., 2009; Rice and Hostert, 1993; Rice and Salt, 1988, 1990).

### 5.2 Environmental Gradients and the Evolution of Dispersal

The effects of environmental gradients and random dispersal on species range evolution are inter-linked. When dispersal is random, the presence of an environmental gradient makes the random core-to-edge gene flow maladaptive, thereby hindering local adaptation at range margins. Therefore, compared with range expansion in a homogeneous environment, the presence of an environmental gradient reduces range expansion speed. The steeper the gradient, the slower the range expansion speed; see Figure 2a for A = 0. Fixing the environmental gradient at a nonzero constant value, the same effect is observed by increasing the dispersal rate. That is, the higher the dispersal, the stronger the effect of maladaptive gene flow, and the slower the range expansion speed (Benning et al., 2024; Shirani and Miller, 2022). Although this observation is counterintuitive, as it may be expected that higher dispersal should increase expansion speed, it can be understood by the fact that the steepness of the environmental gradient and the rate of dispersal are linked through a rescaling of the geographic space. In fact, when dispersal is only random (A = 0 in our model), a change 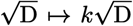 in the dispersal coefficient in the equation of our model (10)–(12) can be absorbed by a rescaling *x* → *x/k* of the space variable, and consequently an equivalent change ∇_*x*_Q → *k*∇_*x*_Q in the environmental gradient (Shirani and Miller, 2022: Remark 3).

In the presence of an environmental gradient, the gene flow created by random dispersal increases local phenotypic variation. The steeper the gradient the higher the level of phenotypic variance; see Figure 2a for A = 0. Our results show that, in extremely steep gradients, inflated phenotypic variation substantially intensifies the phenotypic load that selection imposes on a population’s growth rate. As a result, the population density declines sharply with increases in the environmental gradient; see Figure 2a for A = 0. In particular, the species fails to survive when the gradient exceeds a critical level given by (18). Based on the link we discussed above between the effects of dispersal and environmental gradient, the same pattern of changes in trait variance and population density as shown in Figure 2a (with A = 0) is also observed versus changes in 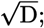 see also Remark 3 by Shirani and Miller (2022). That is, trait variance increase with increases in dispersal, and when dispersal is exceedingly large, population density declines sharply with increases in dispersal. Although the critical gradients we calculated are likely too steep to be plausible for the majority of species (unless possibly representing physical barriers), our results suggest that slowly-growing species under excessively strong selection are quite likely to experience such critical gradients in nature; see Remark 4.2 and Figure 3. This means that, this class of (vulnerable) species are at high risk of extinction in steep environmental gradients if they disperse randomly over long distances.

In light of the link between random dispersal and environmental gradients in hindering local adaption and reducing expansion speed, it is argued that dispersal is strongly selected against in presence of an environmental gradient (Holt, 1985). Using an individual-based model that allows for the evolution of both a fitness-related (niche) trait and a trait that controls dispersal ability, Benning et al. (2024) have shown that the presence of an environmental gradient opposes the evolution of increased dispersal during invasions. If the gradient is steep, it can even result in the evolution of reduced dispersal. This in fact follows from the link between dispersal and environmental gradients we discussed above. Reduced random dispersal is equivalent to a reduced environmental gradient, and hence may be expected to cancel out the effects of steep gradients. However, this conclusion relies on an important assumption, that dispersal evolves as random movements. Although there are stochastic and extrinsic forces that influence dispersal (Lowe and McPeek, 2014), it is not very realistic to assume species evolve random dispersal in steep environmental gradients. By contrast, our results suggests that steep environmental gradients indeed play a key role in evolution of increased phenotype-optimal (adaptive) dispersal, which can then effectively compensate for the disruptive effects of the gradients on local adaptation and range expansion. In view of our results and those of Benning et al. (2024), we can hence hypothesize that there may be different thresholds in the steepness of the gradients that promote evolution of different dispersal strategies. Significantly steep gradients beyond a threshold may promote the evolution of optimal (directed) dispersal. Moderately steep gradients below a threshold may instead lead to the evolution of reduced dispersal.

### 5.3 Phenotype-Optimal Dispersal in Nature

Currently, evidence for matching habitat choice in nature is limited. This may be partly because this mode of adaptation has been overlooked due to a primary focus on natural selection (Edelaar and Bolnick, 2019). However, matching habitat choice may truly be rare because of, for example, high cost for its evolution, inability of species’ individuals to obtain information about their performance and matching habitats, dependence of adaptation on multiple uncorrelated traits, and movement restrictions imposed by strong territoriality (Edelaar et al., 2017; Nicolaus and Edelaar, 2018). Our finding that environmental gradients must be steep for phenotype-optimal dispersal to be consequential suggests an additional non-artifactual reason why this dispersal mode has rarely been observed. Further, we note that we did not explicitly include dispersal costs in our model. Such costs should make phenotype-optimal dispersal even less likely to evolve in shallow gradients. Yet we should note that our estimates of typical values for steepness of environmental gradients are based on limited data (Shirani and Miller, 2022: Sect. 3.2). This is because environmental trait optima and their gradient are hard to measure, and the available measurements are often based on different choices of units which make steepness comparisons almost impossible. Our particular choices of units (see Section 2.5) can provide sufficient generality for future measurements, based on which better estimates for typical steepness of gradients can be obtained.

The current scarcity of evidence for matching habitat choice may also be because efforts for finding such evidence have not been focused on the most promising taxa. As we discussed before, our results suggests that phenotype-optimal dispersal should be more prevalent among highly-mobile slowly-growing species which are possibly exposed to sharp, strong, and frequent changes in their environment. Based on this observation, nomadic birds (Benkman, 2017) and long-range dispersing mammals such as seals, cougars, elephants, buffaloes, rhinoceros, giraffes, and wolves could be examples of species that might specifically benefit from evolving phenotype-optimal dispersal. Microorganisms that live in high chemical or temporal gradients may also optimally move to better matching environment through taxis.

Matching habitat choice is not straightforward to detect in nature. Our results could suggest two preliminary considerations for planning of studies that aim to detect or rule out phenotype-optimal dispersal in the field or laboratory; see, for example, (Camacho et al., 2020; Edelaar and Bolnick, 2012). First, the impacts of gene flow on the fitness of individuals should specifically be observed at population’s range margins. As we discussed before, phenotype-optimal dispersal results in adaptive gene flow to range margins, which is otherwise expected to be maladaptive when dispersal is predominantly random. Second, the level of trait variation should be observed at the core of the population. Phenotype-optimal dispersal can reduce trait variation to strikingly low levels that are unlikely to be maintained by realistically strong selection.

Finally, our results provide a cautionary note. The microcosms (Jacob et al., 2017) or micro-climatic mosaic arenas (Karpestam et al., 2012) that are usually constructed in experiments to detect directed dispersal behavior of a model species may have considerably steeper environmental gradients than those likely to prevail in the species’ natural habitat. Since our results identify a steep environmental gradient as a key factor for evolution of matching habitat choice, we note that researchers should ensure (when possible) that the difference between experimental and natural habitat gradients is insignificant, or use caution in interpreting the experimental results.

### 5.4 Future Research Directions

It is important to note that phenotype-optimal dispersal, the type of matching habitat choice modeled in the present work, incorporates only local environment sensing (assessment). Nonlocal environment sensing, in which an individual can assess the environment at some distance from its location, could readily be incorporated into extensions of our model. Such extensions would be worth studying in part because nonlocal sensing will reduce individuals’ exploratory movements and will increase the chance of settling in globally optimal habitat location. It may be plausible that with nonlocal sensing, the effects of matching habitat choice we have identified here would be significant over shallower environmental gradients. The present study suggests that with purely local resource sensing, matching habitat choice should not be a favored adaptive mechanism in many environments. Since we have performed all our studies in a linear environment, in which local and global sensing will have the same outcomes, we still suspect that our hypothesis remains valid even in case where individuals have nonlocal assessment capabilities. Studies with nonlocal sensing, possibly with nonlinear habitat gradients, would help test the robustness of our hypothesis more rigorously.

Phenotypic adaptation to a heterogeneous environment can occur through several modes, including habitat choice, natural selection, and phenotypic plasticity. How strong each of these modes is relative to the others, how they interact, and under what conditions they are more favored are questions that are not yet fully addressed (Edelaar and Bolnick, 2019; Edelaar et al., 2017; Scheiner et al., 2022). We included phenotype-optimal dispersal and selection in our model. We observed that strong optimal dispersal tends to reduce trait variation relatively fast, leaving less genetic variation for selection to act upon. When a population experiences a sharp change in the environment, it suffers from a strong selective load. We observed that optimal dispersal is quite effective in mitigating the effects of this load and facilitating rapid recovery. This implies that, in rapidly changing environments, natural selection can indeed increase the potential for evolution of optimal dispersal. Detailed analysis of the interactions between natural selection and matching habitat choice, and further extensions to include other modes such as plasticity, are important directions for future work. The extensive literature on the evolution of phenotypic plasticity and its effects on range dynamics (Agrawal, 2001; Chevin and Hoffmann, 2017; Chevin and Lande, 2011; Chevin et al., 2010; DeWitt and Scheiner, 2004; DeWitt et al., 1998; Eriksson and Rafajlović, 2022; Forsman, 2015; Ghalambor et al., 2007; Murren et al., 2015; Pigliucci, 2005; Scheiner, 1993; Scheiner et al., 2012; Schlichting and Pigliucci, 1998; Schmid et al., 2019; Turko and Rossi, 2022; Valladares et al., 2014) as well as the preliminary results on coevolution and comparison of phenotypic plasticity with matching habitat choice (Boyle and Start, 2020; Edelaar et al., 2017; Lowe and Addis, 2019; Nicolaus and Edelaar, 2018; Scheiner, 2016) can help guide this topic of research.

## Declarations

- Funding: not applicable
- Conflict of interest/Competing interests: The authors declare no conflict of interest.
- Ethics approval: not applicable
- Availability of data and materials: This manuscript has no associated data

## Appendix A Perceived Environmental Gradient

The perceived gradient partially defined in (9) over a smaller habitat Ω_*δ*_ requires certain technical considerations near the boundary of the habitat so that it can be extended to the whole habitat Ω. To include such an extension, we assume that the individuals of the species are able to perceive the boundary of the habitat once they become sufficiently close to it (closer than the constant *δ*,) and they avoid crossing the boundary. This means that, we assume that the normal component of 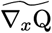 to the boundary of Ω is zero. Over a neighborhood of width *δ* around the boundary, we smoothly extend the 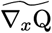 defined by (9) so that its normal component to the boundary gradually vanishes to zero. As a result, the extended 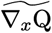 will be tangential to the boundary of Ω. For this, we first define the following cut-off function

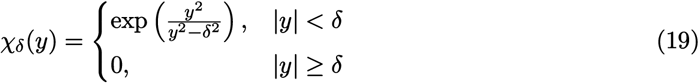

which smoothly declines from 1 at *y* = 0 to 0 at |*y*| = *δ*.

Now, for a rectangular habitat Ω = (*a*_1_, *b*_1_) × … × (*a*_m_, *b*_m_), we extend (9) as

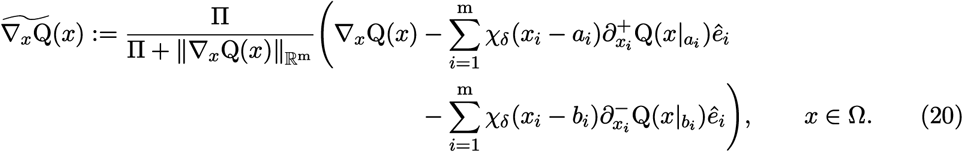

where *ê*_*i*_ denotes the *i*th standard unit vector in ℝ^m^, and 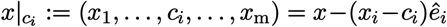. Moreover, 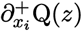 and 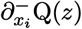 denote, respectively, the right-hand and left-hand partial derivatives of Q with respect to *x*_*i*_ evaluated at a point *z*. The summation terms in (20) gradually remove the normal component of ∇_*x*_Q to the boundaries *x*_*i*_ = *a*_*i*_ and *x*_*i*_ = *b*_*i*_, over a *δ*-neighborhood of the boundaries, so that 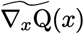 eventually becomes completely tangent to the entire boundary of Ω. The removal of the normal components of the perceived gradient 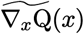 to habitat boundary automatically results in no phenotype flux through the boundary due to the directed dispersal term (1b). Therefore, the no-flux (reflecting) boundary conditions that we discussed in Remark 1 and Appendix A.5 of our previous work (Shirani and Miller, 2022) can also be applied to the model (10)–(12) we present in this work. If a periodic boundary condition is considered across the *j*th spatial direction, then the terms associated with *i* = *j* are excluded from the summation terms in (20). This is because imposing periodic boundary condition in a spatial direction is equivalent to considering a habitat that is periodically extended in that direction.

For the one-dimensional habitats Ω = (*a, b*) used in the results presented in Figures 1–7, the perceived environmental gradient (20) can be written as

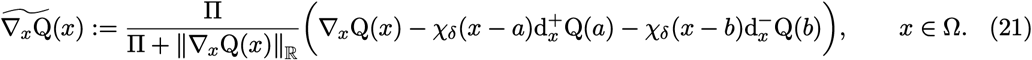

where 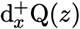 and 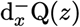 denote, respectively, the right-hand and left-hand derivatives of Q with respect to *x* evaluated at a point *z*. Note that ∥∇_*x*_Q(*x*)∥_ℝ_ = |d_*x*_Q(*x*)|.

For the two-dimensional habitat Ω = (*a*_1_, *b*_1_) ×(*a*_2_, *b*_2_) used in the results presented in Figure 8, we considered reflecting boundary conditions at *x*_1_ = *a*_1_ and *x*_1_ = *b*_1_, and periodic boundary conditions across the *x*_2_-axis. Therefore, the perceived environmental gradient for this problem can be written as

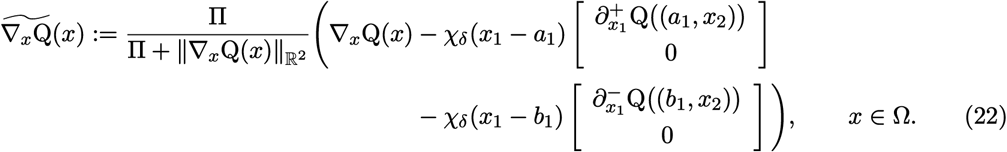

In all of the numerical simulations presented in this work, we set the maximum perceived gradient to be Π = 1 Q/X, and we assume the individuals can sense the habitat boundary at a distance smaller than or equal to *δ* = 2 X.

## Appendix B Model Derivation

We use the basic equation (1) to derive the equations of our model (10)–(12). As the constructing components of the basic equation (1), we use the intrinsic growth rate (5), the perceived dispersal force (8) which substitutes for −∇_*x*_*θ* in (1b), and the rate of mutational changes 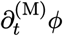 given by Equation (20) in our previous work (Shirani and Miller, 2022). Our derivation relies on the major assumptions (i)–(viii) provided in Section 2.1.

The basic equation (1) that we use to derive the equations of our model differs from the the Equation (25) in our previous work (Shirani and Miller, 2022) only in the inclusion of the optimal dispersal term (1b). As a result, applying the derivation steps that we perform below to the rest of the terms, (1a), (1c), and (1d), will give the terms (10a), (10c), (11a), (11b), (11e), (11f), (12a)–(12c), and (12f), whose derivation can be obtained as a single-species version of our general multi-species model presented in our previous work; see, for example, the Equation (12)–(14) given in that work for a one-dimensional habitat. Therefore, we exclude the derivation of these terms from our work here, and refer the reader to our previous work for details of those derivations. In the following, we show the derivation of the new terms (10b), (11c)–(11e), (12d) and (12e) that model the effects of optimal dispersal.

We first substitute (8) into the basic equation (1) to write

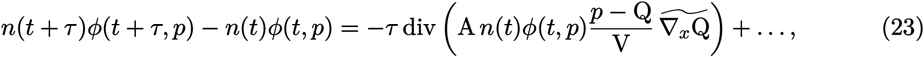

where “… “denotes the terms we have excluded from our derivation, as stated above. Note that, for simplicity of exposition, in writing (23) and the rest of the derivations that follow, we do not explicitly show the dependence of the variables and parameters on *x*.

Now, we integrate both sides of (23) with respect to *p* over ℝ to obtain

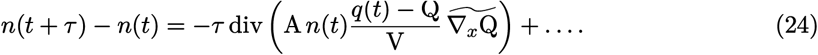

Dividing both sides of (24) by *τ* and taking the limit as *τ* → 0 yields (10).

To derive (11), we multiply both sides of (23) by *p* and integrate the result with respect to *p* over R. Noting that ∫_ℝ_*p*^2^*φ*(*t, p*)d*p* = *v*(*t*) + *q*^2^(*t*), we obtain

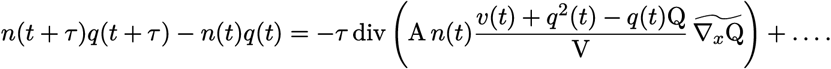

which, after dividing by *τ* and taking the limit as *τ* → 0, gives

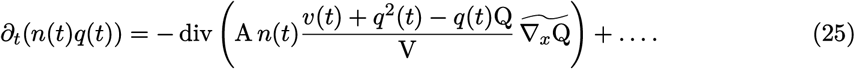

Now, we use the product rule on the left-hand side of (25) and substitute (10) into the result to obtain

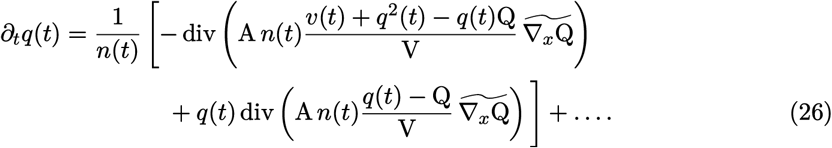

Denoting the first term within the brackets in (26) by [(26).1st], we can write

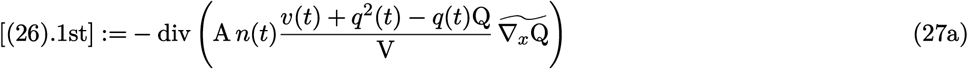

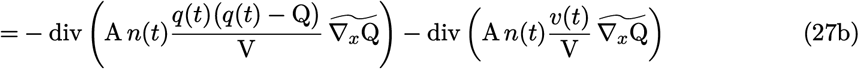

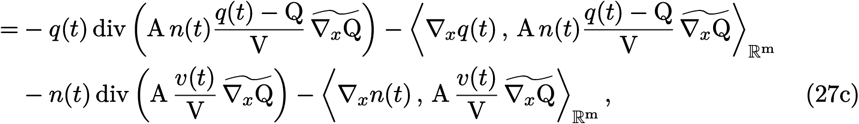

where the product rule for divergence of a scalar field times a vector field has been applied to each of the two divergence terms of (27b) to obtain (27c). For this, *q*(*t*) has been considered as the scalar field in the first divergence term in (27b), and *n*(*t*) has been considered as the scalar field in the second divergence term. Now, we substitute the result into (26) for the first term within the brackets and obtain (11).

Finally, to derive (12), we first note that

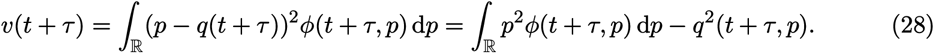

We multiply both sides of (28) by *n*(*t* + *τ*) and use (23) to write

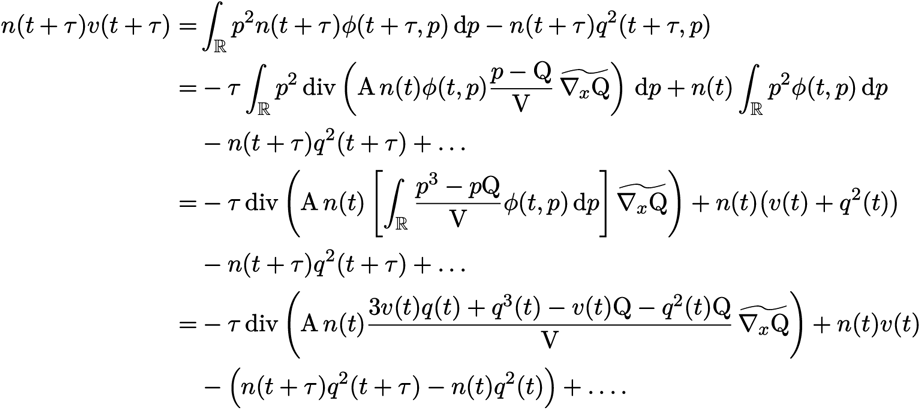

Next, we subtract *n*(*t*)*v*(*t*) from both sides of the above equation, divide both sides by *τ*, and take the limit as *τ* → 0. We obtain

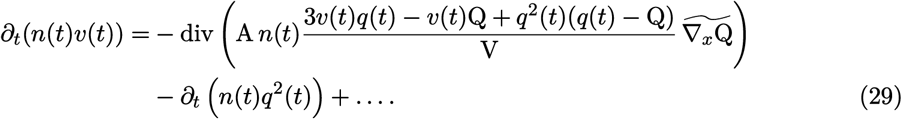

If we use the product rule to write *∂*_*t*_(*n*(*t*)*v*(*t*)) = *n*(*t*)*∂*_*t*_*v*(*t*) + *v*(*t*)*∂*_*t*_*n*(*t*), wherein *∂*_*t*_*n*(*t*) is given by (10), and split the divergence term in (29) into three terms, we can write

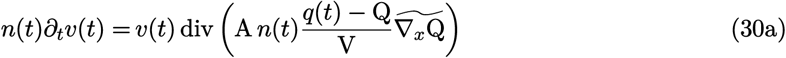

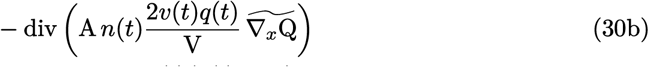

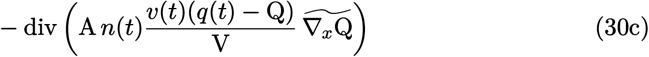

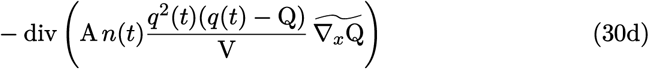

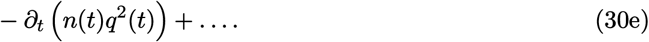

Note that, in the derivation of (30), the terms involving (10a) and (10c) are absorbed into the “…” in (30e). This is because these terms are not associated with the model of phenotype-optimal dispersal and, as we stated above at the beginning of this section, they are handled in the derivations following the same steps taken in our previous work.

Now, for (30b), we use the product rule for divergence of a scalar field times a vector field, with the scalar field *n*(*t*)*q*(*t*), along with the product rule for ∇_*x*_(*n*(*t*)*q*(*t*)), to write

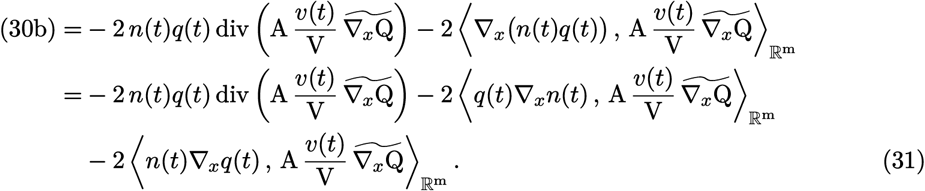

Similarly, applying the product rule for the divergence in (30d), keeping in mind *q*^2^(*t*) as the scalar field, and noting ∇_*x*_*q*^2^(*t*) = 2*q*(*t*)∇_*x*_*q*(*t*), we have

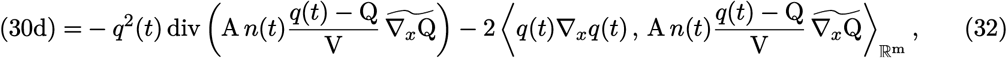

For (30e), we use (10) and (11) to write

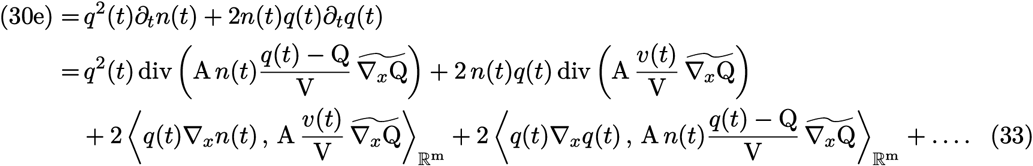

Note that, in writing the first inner product term in (33), we used *n*(*t*)∇_*x*_ log *n*(*t*) = ∇_*x*_ log *n*(*t*)). Moreover, similar to what we noted above, we have absorbed into the “… “in (33) all those terms that do not correspond to the model of phenotype-optimal dispersal in (10) and (11).

Finally, denoting the right-hand side of (30a) by (30a).RHS and using the product rule for divergence (with the scalar field *v*(*t*)), we can write

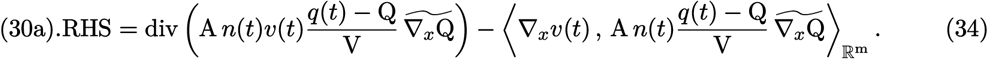

We then obtain (12) by substituting (31), (32), (33), and (34) into (30) and simplifying the result. This completes the derivation of the model.

---

**Figure S1:**
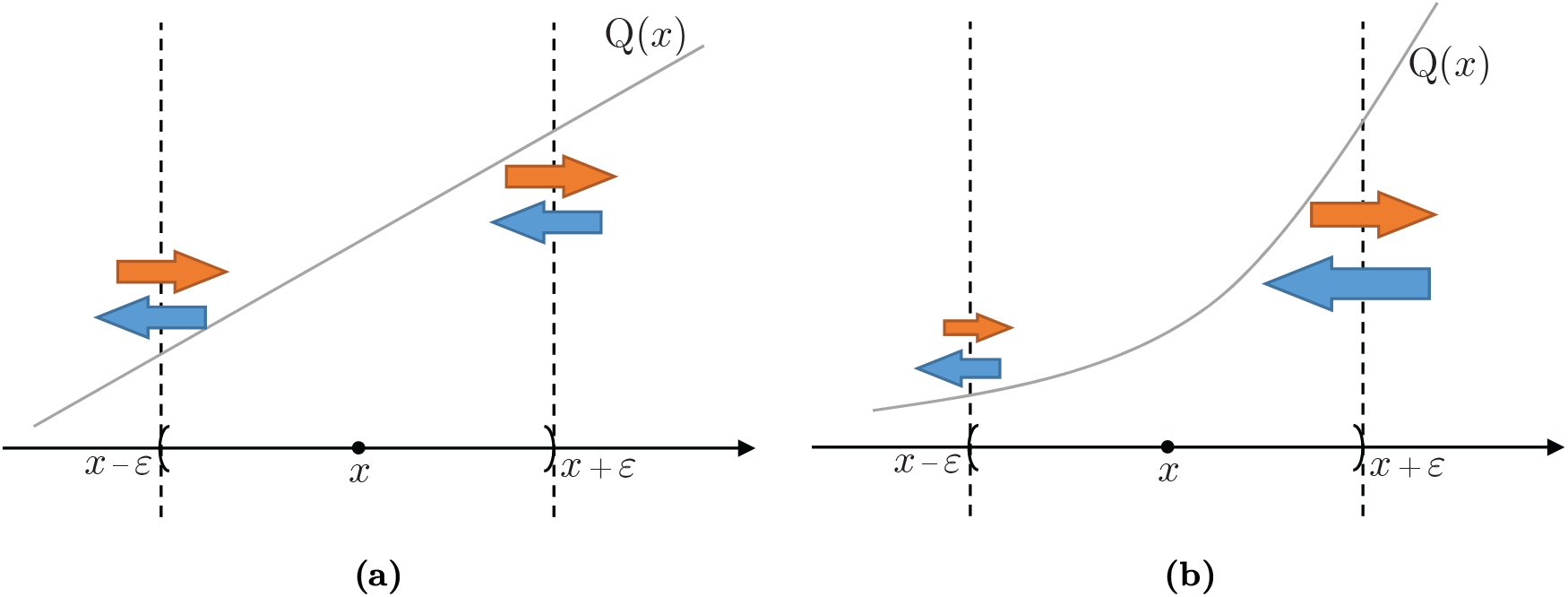
Effects of divergence in (perceived) environmental gradient on the mean trait value of a local population. We consider a local population located at a small neighborhood (*x* − *ε, x* + *ε*) of a habitat point *x*. To illustrate changes in the trait mean that are purely caused by divergence in the perceived environmental gradient, that is the effects of the term (14b) in Equation (14) of the main text, we assume that the population density *n*, the trait variance *v*, and the optimal dispersal propensity A take constant values over a sufficiently large neighborhood of *x*, including (*x* − *ε, x* + *ε*). Moreover, we assume that the dispersal rate of the individuals is independent of the magnitude of their phenotype-environment mismatch. Only the sign of the mismatch will determine whether they move in the direction or in the opposite direction of the environmental gradient. For simplicity, we further assume that the perceived gradient 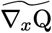 is equal to the actual gradient ∇_*x*_Q. As a result, the directed (optimal) dispersal rate will only be controlled by the magnitude of the environmental gradient. In each graph, the flow of individuals with higher phenotype value into and out of the *ε*-neighborhood is shown by red arrows. The blue arrows show the flow of lower phenotype values. Larger arrows indicate higher dispersal (flow) rates. The trait optimum Q(*x*) is increasing in *x*. As a result, the internal individuals that choose to move out of the neighborhood through the right boundary have larger phenotype values than the external individuals that choose to move into the neighborhood through this boundary. The opposite holds at the left boundary. In (a), the environmental gradient is constant, and we have div 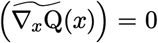. As a result, the inward and outward dispersal rates are equal at both boundaries, and their net effect does not change the mean trait value of the local population. In (b), the environmental gradient is increasing, and we have div 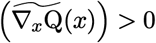. As a result, the internal individuals that move out through the right boundary perceive a lower environmental gradient than the gradient perceived by external individuals that move in. That means, the outward dispersal rate at the right boundary is lower than the inward rate. The opposite holds at the left boundary. Moreover, since the environmental gradient at the right boundary is higher than the gradient at the lower boundary, the overall dispersal rate at the right boundary is higher than the rate at the left boundary. Consequently, as the arrows indicate, the local population loses more of its larger-valued phenotypes than it gains, and receives more of smaller-valued phenotypes than it loses. This leads to a reduction in the mean trait value of the local population due to the positive divergence in the environmental gradient.

**Figure S2:**
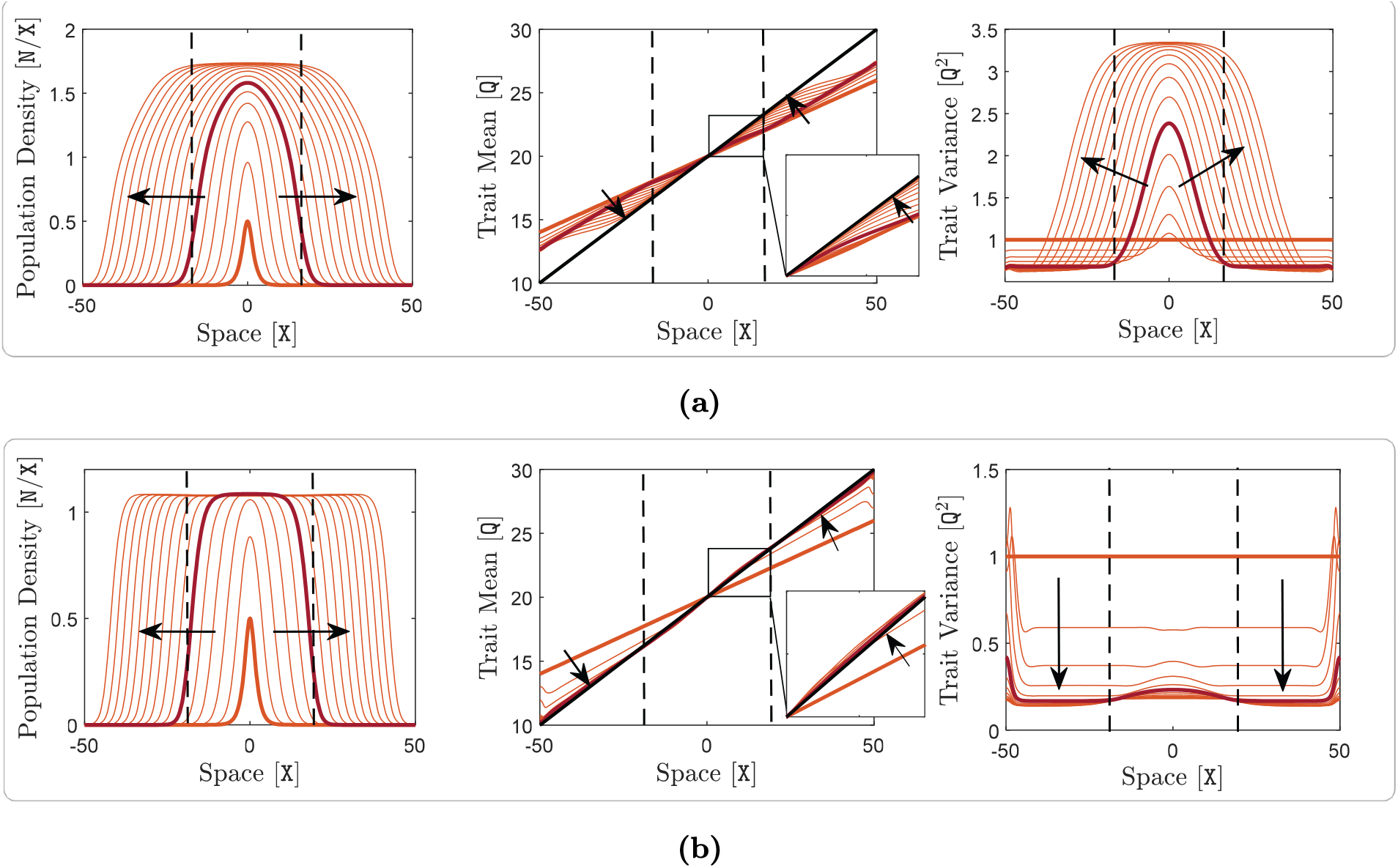
Adaptive range dynamics of a species with specialist individuals in a one-dimensional habitat. Here, m = 1 and A(*x*) is constant, taking different values in panels (a) and (b). The phenotype utilization variance takes the small value of V = 1 Q^2^. The rest of the model parameters, including the environmental gradient, take their typical values given in Table 1 of of the main text. Panel (a) shows the range expansion dynamics of a species without optimal dispersal, A = 0 X^2^/T, whereas panel (b) shows the range expansion dynamics of a species with strong optimal dispersal, A = 10 X^2^/T. In all graphs, including the insets of trait mean graphs, curves are shown at every 1 T and a sample curve at *t* = 5 T is highlighted in red. The same description as given in Figure 1 of the main text holds here for the quantities shown in each panel, the thick curves, arrows, and dashed lines.

**Figure S3:**
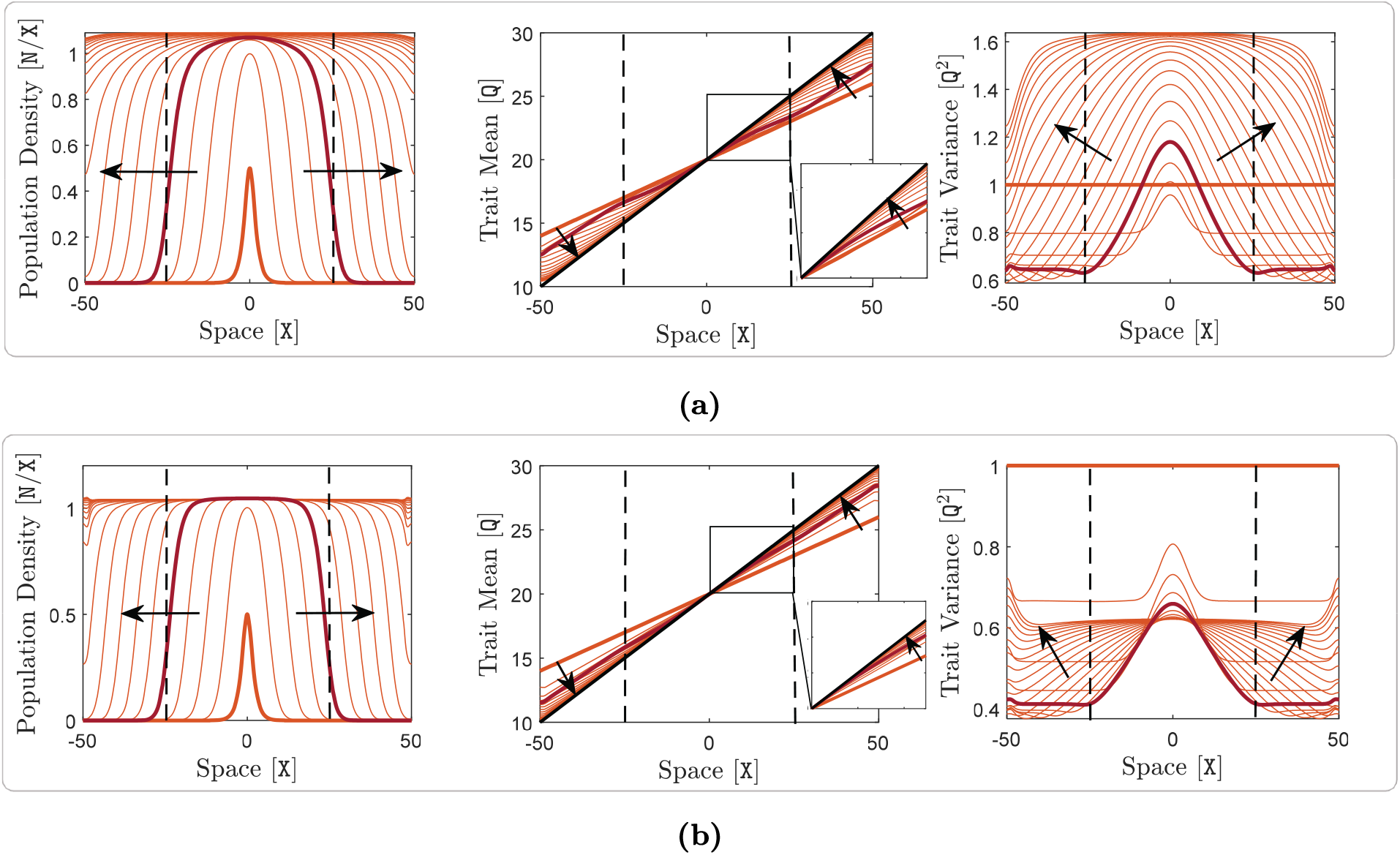
Adaptive range dynamics of a species in a one-dimensional habitat with typical (shallow) environmental gradient. Here, m = 1, Q(*x*) is linear with the typical gradient of ∇_*x*_Q = 0.2 Q/X, and A(*x*) is constant, taking different values in panels (a) and (b). The rest of the model parameters take their typical values given in Table 1 of the main text. Panel (a) shows the range expansion dynamics of a species without optimal dispersal, A = 0 X^2^/T, whereas panel (b) shows the range expansion dynamics of a species with strong optimal dispersal, A = 10 X^2^/T. In all graphs, including the insets of trait mean graphs, curves are shown at every 2 T. The same description as given in Figure 1 of the main text holds here for the curves, arrows, and dashed lines.

**Figure S4:**
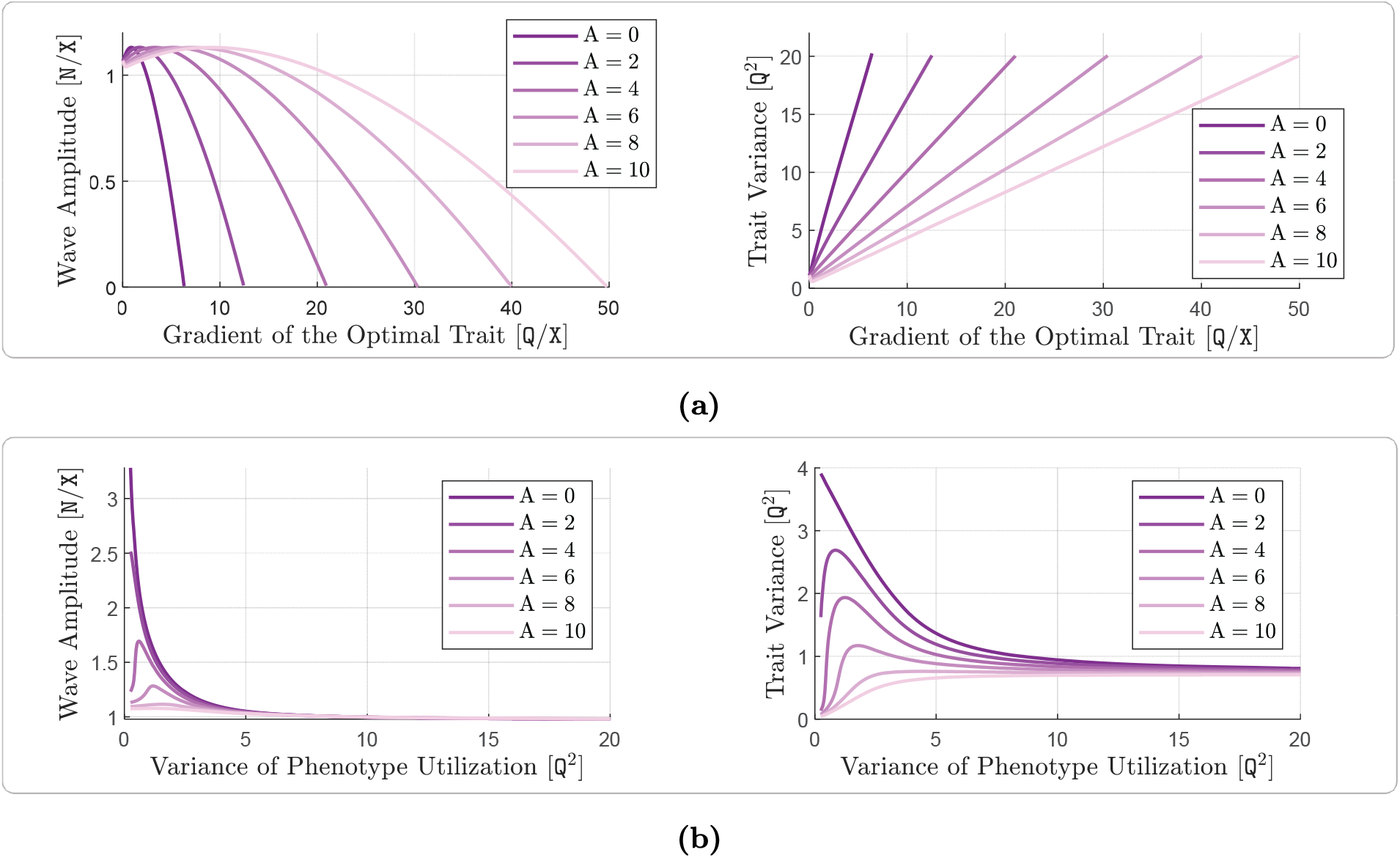
Effects of optimal dispersal on range expansion waves and maximum intraspecific trait variance of a species. The graphs shown in this figure directly correspond to those shown in Figure 2 of the manuscript, with the only difference being that here the curves are computed more accurately based on the spatially homogeneous equilibrium of the model; see our previous work for details of such analysis (Shirani and Miller, 2022: Sect. 4.2). Specifically, m = 1, and A(*x*) takes six different constant values in the range from 0 X^2^/T to 10 X^2^/T. The trait optimum Q(*x*) is linear in *x*, with variable gradient in panel (a) and the constant gradient of ∇_*x*_Q = 0.2 Q/X in panel (b). The phenotype utilization variance takes the constant value V = 4 Q^2^ in panel (a), and is variable in panel (b). The rest of the model parameters take their typical values given in Table 1 of the main text. In each panel, variations in the amplitude of the traveling waves are shown on the left and variations in the maximum intraspecific trait variance is shown on the right. Panel (a) shows the effects of different levels of optimal dispersal at different values of the environmental gradient ∇_*x*_Q. Panel (b) shows the effects of different levels of optimal dispersal at different values of the individuals’ phenotype utilization variance V.

**Figure S5:**
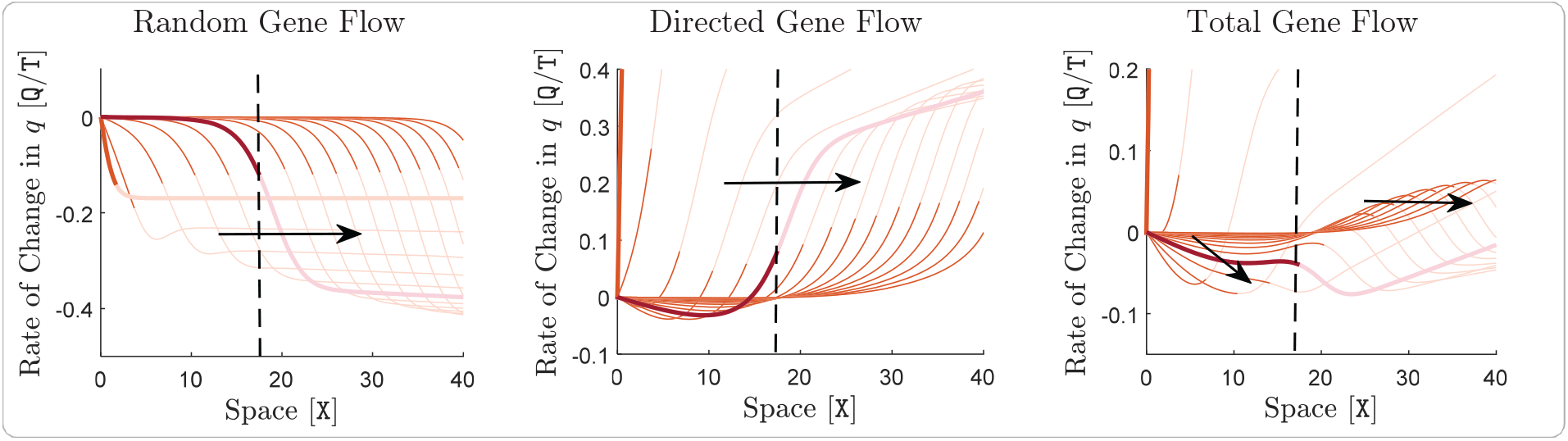
Effects of gene flow on local adaption of a species. Contribution of random gene flow to the rate of change of the trait mean, *∂*_*t*_*q*, is shown on the left, contribution of directed gene flow generated by optimal dispersal is shown in the middle, and contribution of the total gene flow is shown on the right. The curves in each graph are calculated using the components of the trait mean equation as described in Figure 3 of the main text. The graphs correspond to the same simulation of species’ range dynamics given in Figure S2b, that is, when V = 1 Q^2^ and A = 10 X^2^/T. In all graphs, the evolution of the computed curves are shown only on the right half of the habitat. The curves extend symmetrically about the origin to the left half of the habitat. Moreover, the portion of each curve that lies outside the effective range of the species, that means over the regions where the population density rapidly vanishes to zero, has been made transparent. The descriptions of the time increments, highlighted curves, arrows, and dashed lines are the same as those provided in Figure S2b.

**Figure S6:**
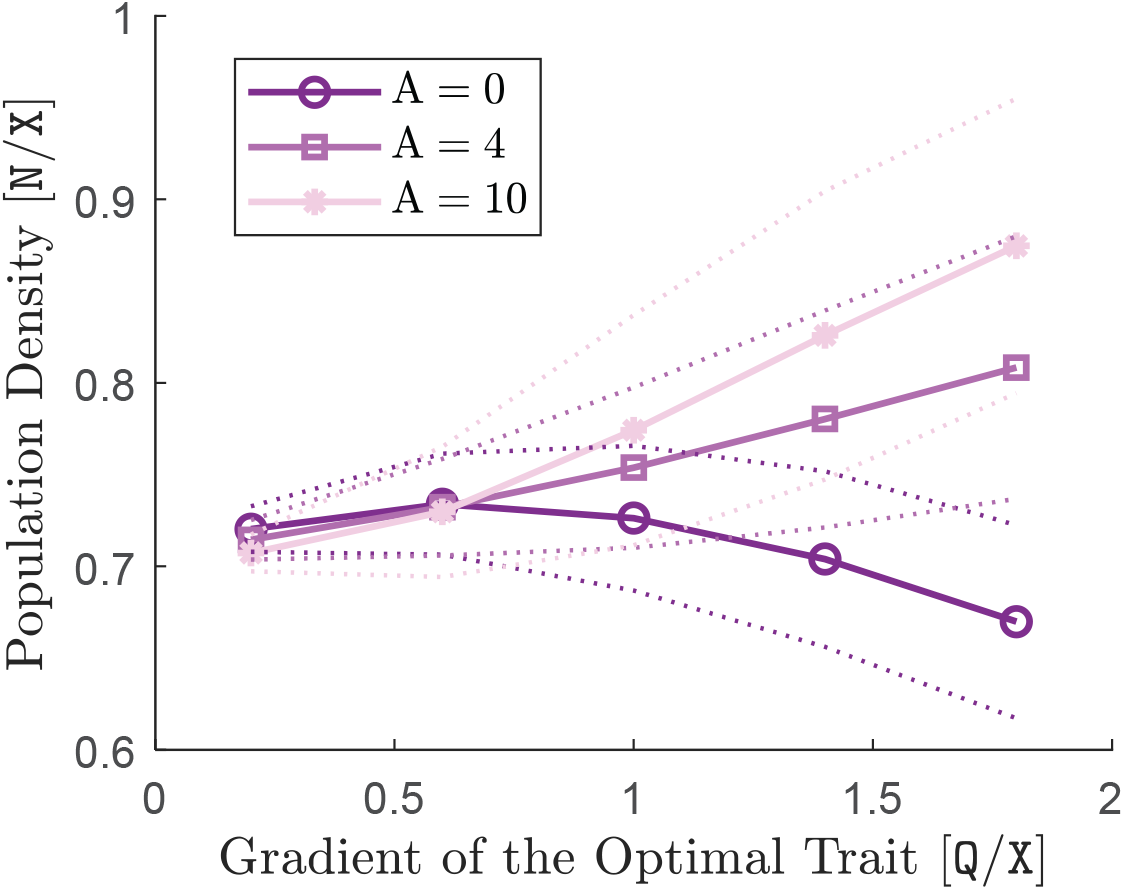
Steady-state mean population density of a species under periodic abrupt fluctuations in the environmental trait optimum. The simulation setup is the same as that of Figure 5a in the main text, with the only difference being that the gradient of the optimal trait and A are made variable here. At each value of the gradient and each value of A, the simulation is run for a sufficiently long period of time so that the amplitude of fluctuations in the population density reaches a steady-state. The minimum and maximum values of such periodic fluctuations (peaks of the blue and red curves as shown in Figures 5a and 5b in the main text) are calculated near the end of the simulation, and are plotted by (interpolated) dotted lines. The average value of the fluctuations is shown by different markers for each value of A, along with an interpolated solid line.

**Figure S7:**
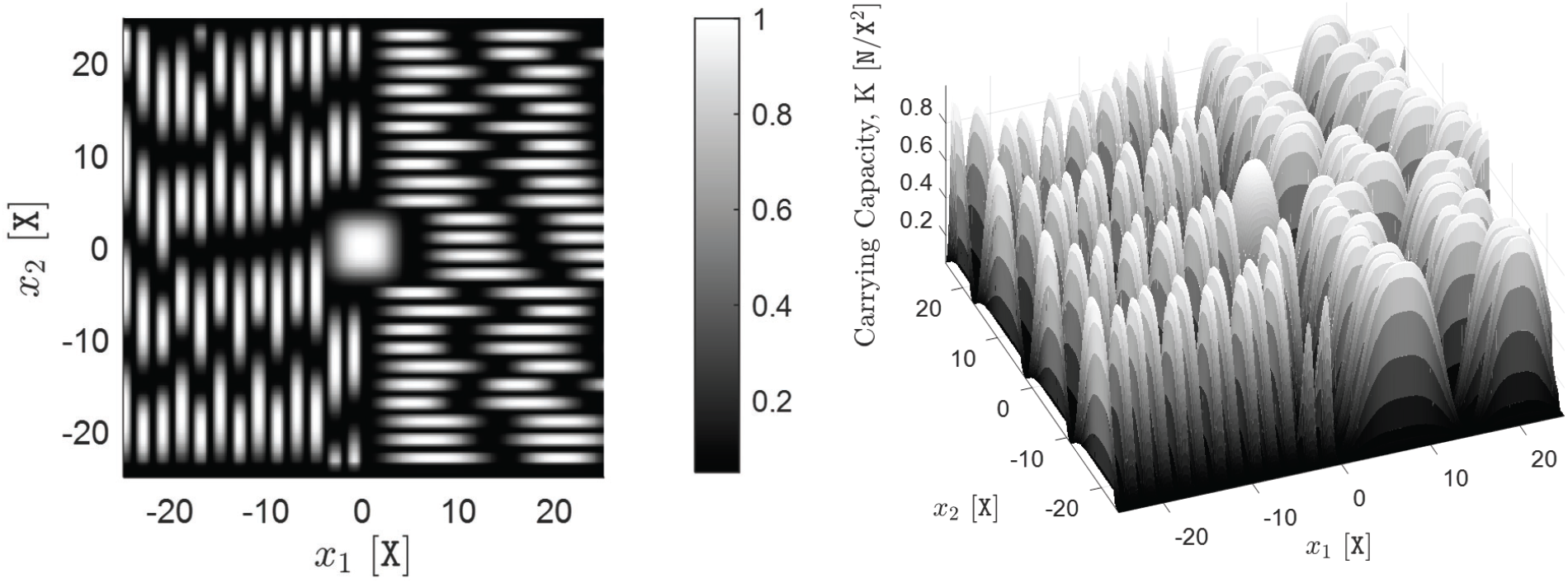
Carrying capacity of the two-dimensional fragmented habitat. The spatial pattern of the carrying capacity K(*x*) used in the simulation of the two-dimensional fragmented habitat of Figure 7 in the main text. The parameter K takes its maximum value of 1 N/X^2^ at the center of each patch, and its minimum value of 0.05 N/X^2^ at the borderlines between the patches.

**Figure S8:**
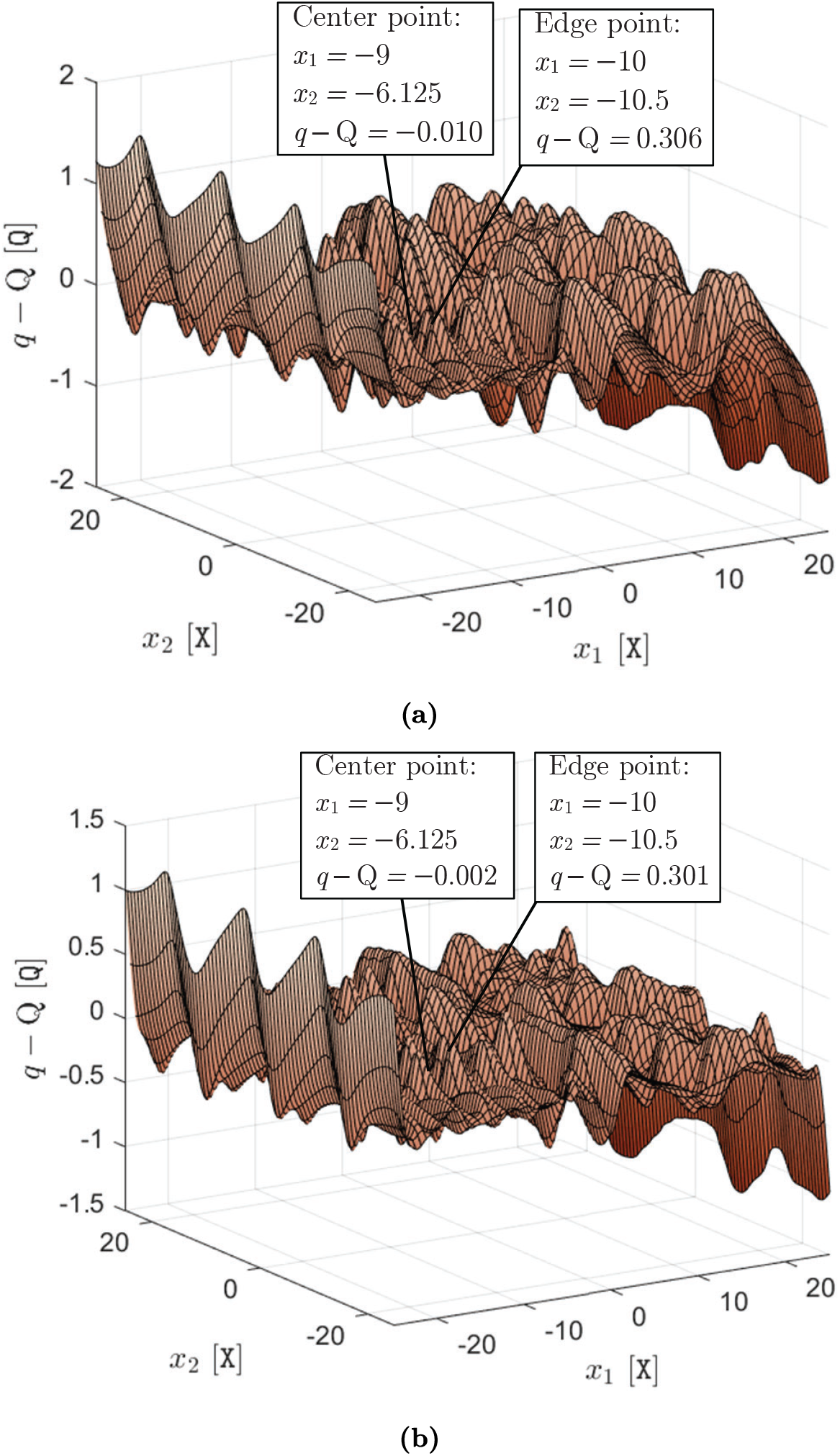
Phenotype-environment mismatch of a species in a fragmented habitat. The results shown here correspond to the same simulation results shown in Figure 7 of the main text, for a species that spreads over a fragmented two-dimensional habitat. In each graph, the approximate steady-state profile of the phenotype-environment mismatch *q* − Q obtained at *t* = 40 T is shown. Graph (a) shows the mismatch for the species with A = 0 X^2^/T (upper panel of Figure 7), and graph shows the mismatch for the species with A = 10 X^2^/T (lower panel of Figure 7). Two samples of the exact values of the mismatch, one at the center of a patch and the other at its edge, are also explicitly specified in the graphs.

**Figure S9:**
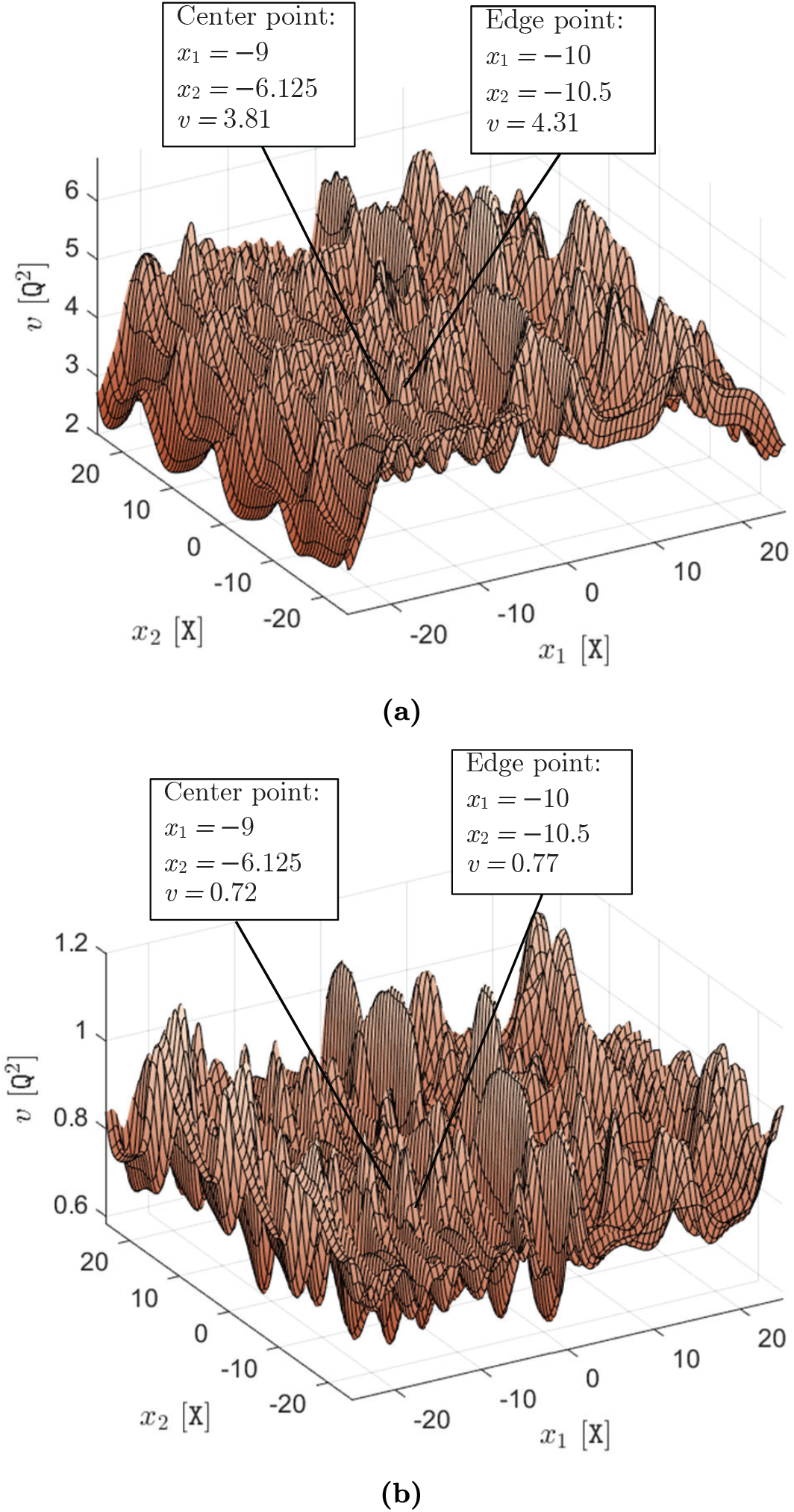
Intraspecific trait variance of a species in a fragmented habitat. The results shown here correspond to the same simulation results shown in Figure 7 of the main text, for a species that spreads over a fragmented two-dimensional habitat. In each graph, the approximate steady-state profile of the intraspecific trait variance *v* obtained at *t* = 40 T is shown. Graph (a) shows the trait variance for the species with A = 0 X^2^/T (upper panel of Figure 7), and graph (b) shows the trait variance for the species with A = 10 X^2^/T (lower panel of Figure 7). Two samples of the exact values of the trait variance, one at the center of a patch and the other at its edge, are also explicitly specified in the graphs.

## Footnotes

1 We use the term “intrinsic” growth rate for the logistic growth rate *g*(*x, t, p*), given by (5), to distinguish this component of the population’s instantaneous growth rate in our model from the other components contributed by dispersal and mutations. This makes our interpretation of the intrinsic growth rate slightly different from an alternative definition that specifies it as the growth rate of the population as its density goes to zero. In our model, this alternative definition is specified as the “maximum” intrinsic growth rate, denoted by the parameter R in (5).

2 We note that *ψ*_*p*_ itself (instead of its first-order approximation) does not serve as a fully meaningful scalar potential energy function. Setting 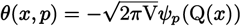 would induce a dispersal force −∇_*x*_*θ* as 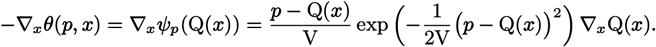 Therefore, when phenotype-environment mismatch |*p* − Q(*x*) → ∞, the dispersal force −∇_*x*_*θ*(*p, x*) → 0, which does not make sense.

3 Note that, as stated in Assumption (vi) and described in Remark 2.2, in writing the phenotype utilization distribution (3) we assume an identification between the resource axis and the phenotype axis.

4 We recall the product rule for divergence of a scalar field times a vector field as 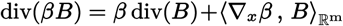, where *β* denotes a scalar field, whereas *B* denotes a vector field.

5 The Laplacian Δ*f* (*x*) := div(∇_*x*_*f* (*x*)) of a function *f* at a point *x* is essentially equal to the average of the differences *f* (*y*) − *f* (*x*) when *y* takes all values over a small ball ℬ(*x*) centered at *x*. In other words, Δ*f* (*x*) essentially gives the amount by which the average value of *f* over ℬ(*x*) differs from *f* (*x*).

